# A neural network model of free recall learns multiple memory strategies

**DOI:** 10.1101/2025.09.25.678592

**Authors:** Moufan Li, Kristopher T. Jensen, Qiong Zhang, Qihong Lu, Marcelo G. Mattar

**Affiliations:** Department of Psychology, New York University; Sainsbury Wellcome Centre, University College London; Department of Psychology, Rutgers University—New Brunswick; Department of Computer Science, Rutgers University—New Brunswick; Department of Neuroscience, City University of Hong Kong; Center for Neural Science, New York University; Tandon School of Engineering, New York University

## Abstract

Humans exhibit structured patterns of memory recall, including a tendency to recall more recent information and to recall events in the same order they were experienced. Classic computational models explain these patterns by positing that memories incorporate the ongoing “temporal context”, formed by smoothly integrating the stimulus history. However, it is unclear whether a single mechanism can account for the full repertoire of human memory strategies, as the optimal approach may be task-dependent. For example, human memory experts widely apply the “memory palace” strategy, which is empirically better but not captured by temporal context models. Here we show that neural networks optimized for free recall develop diverse retrieval strategies, with only some of them resembling temporal context models. The best-performing models discovered a stimulus-invariant index code that emphasizes the studied position of each list item, instead of its temporal context. This creates a stable scaffold for forward recall akin to the memory palace technique. This index code was more likely to emerge when networks were i) encouraged to recall all studied items rather than prioritizing only the next few items, and ii) prevented from relying on recency. Our findings demonstrate that human-like recall patterns can arise from multiple distinct computational mechanisms, and that sequential retrieval using item index is an optimal strategy that explains expert-level recall performance.

## 1 Introduction

Decades of research into episodic memory have sought to understand how humans search their past experiences to recall information (Tulving and Thomson, 1973; Johnson and Rugg, 2007; Polyn and Kahana, 2008). The free recall task, where participants study a list of items and then recall them in any order, has become a central paradigm for this investigation. Human recall in this paradigm consistently reveals highly structured patterns, including serial position effects of enhanced memory for the first and last items (Murdock Jr, 1962), and the temporal contiguity effect, describing people’s tendency to recall memories in nearby list positions to the just recalled memory and in the same order as they were studied (Kahana, 1996; Howard and Kahana, 1999). These patterns, replicated and extended over decades of research (Glenberg et al., 1980; Koppenaal and Glanzer, 1990; Zaromb et al., 2006; Polyn et al., 2009b; Healey et al., 2019), provide strong constraints on theories of how episodic memory organizes and retrieves information. Yet, these general patterns coexist with striking variability in individual memory ability and strategy (Healey et al., 2014; Unsworth, 2019). This raises a crucial question: what computational mechanisms can account for this diversity, and under what circumstances is a particular strategy optimal?

A prominent class of computational models explains these memory patterns by positing the existence of a gradually changing internal state, or temporal context, that evolves with each experienced item and becomes associated with them in memory. The most influential of these, the Temporal Context Model (TCM), proposes that this context is reinstated during memory recall and serves as a dynamic cue to retrieve subsequent memories (Howard and Kahana, 2002; Sederberg et al., 2008; Polyn et al., 2009a). This simple mechanism explains many recall phenomena such as the temporal contiguity effect, and has become the standard model of free recall. However, it is possible that alternative computational strategies, distinct from a drifting temporal context, could also support effective recall performance and account for the observed recall patterns.

One important alternative computational strategy for free recall is the “memory palace” (method of loci), a technique widely used by memory experts to achieve extraordinary recall performance (Roediger, 1980; Wilding and Valentine, 1997). The memory palace strategy involves mentally placing items along a sequence of imagined familiar locations during encoding and systematically retracing these imagined locations during retrieval (Maguire et al., 2003; Dresler et al., 2017; Wagner et al., 2021). Computationally, this approach is fundamentally different from TCM. Instead of relying on a drifting, content-based context, the memory palace creates a stable scaffold by representing each item’s serial position independently of its content—a mechanism more akin to positional coding models (Neath and Crowder, 1990; Brown et al., 2000; Logan and Cox, 2021). Although rational analysis (Anderson and Milson, 1989; Lieder and Griffiths, 2020) suggests that such forward-ordered recall is indeed an optimal policy for free recall (Zhang et al., 2023), this work has been constrained by the architectural assumptions of TCM. Critically, these assumptions cannot account for the stable, “index”-based strategy of the memory palace, leaving it unclear how to computationally explain expert-level performance and what alternative optimal strategies might exist. We therefore ask what computational strategy a system trained to maximize performance converges to. Does it rely on a drifting, content-based temporal context, or would does it instead discover a different mechanism, such as an index-based strategy?

Here, we use task-optimized neural networks to discover optimal memory strategies empirically (Richards et al., 2019). This approach provides more flexibility for context updating compared to hand-crafted cognitive models, such as TCM. We designed a model composed of a recurrent neural network (RNN) augmented with an external episodic memory buffer (Lu et al., 2022; Whittington et al., 2020) and trained it using reinforcement learning. The model was optimized with the sole objective of maximizing the number of recalled items, without any constraints on the order of recall. By analyzing the internal representations and behaviors that emerge from this optimization process, we can identify the computational principles that govern effective memory search, thus offering a complementary perspective on the diversity of strategies that can facilitate free recall.

Our results reveal that neural networks trained solely to maximize recall performance could develop various computational strategies, with many of them producing human-like recall patterns such as the temporal contiguity effect. Remarkably, while some networks converged on TCM-like mechanisms based on drifting temporal context, other networks discovered different computational strategies. A particularly common strategy discovered by some networks involved representing each item’s abstract serial position independently of content, and systematically traversing these index representations during retrieval to recall items in strict forward order — a strategy functionally analogous to the memory palace technique. This index-based strategy achieved superior performance and robustness compared to TCM-like mechanisms, providing a normative account for why perfect forward-ordered recall is so effective and supporting theoretical claims about its optimality (Zhang et al., 2023). These findings establish that multiple computational strategies can produce human-like recall, and that learning an item-independent scaffold for memory organization leads to a powerful, and potentially optimal, solution for free recall.

## 2 Results

### 2.1 A neural network model of sequential episodic memory

To investigate the computational principles of memory retrieval in distributed neural system, we simulated the classic free recall task (Figure 1a). This task consists of a *study* phase where a list of items is presented sequentially, followed by a *response* phase where the model must retrieve as many items as possible in any order. We designed a neural network model capable of encoding and retrieving sequential memories (Figure 1b-c). The model consists of a “context module” that uses gated recurrent units (GRUs) (Cho et al., 2014) as working memory, and a “memory module” with slots for storing episodic memories (Ritter et al., 2018; Lu et al., 2022). This architecture, while analogous to the context and feature spaces in the Temporal Context Model (TCM) (Howard and Kahana, 2002), provides additional flexibility allowing for the emergence of novel context-updating mechanisms.

**Figure 1:**
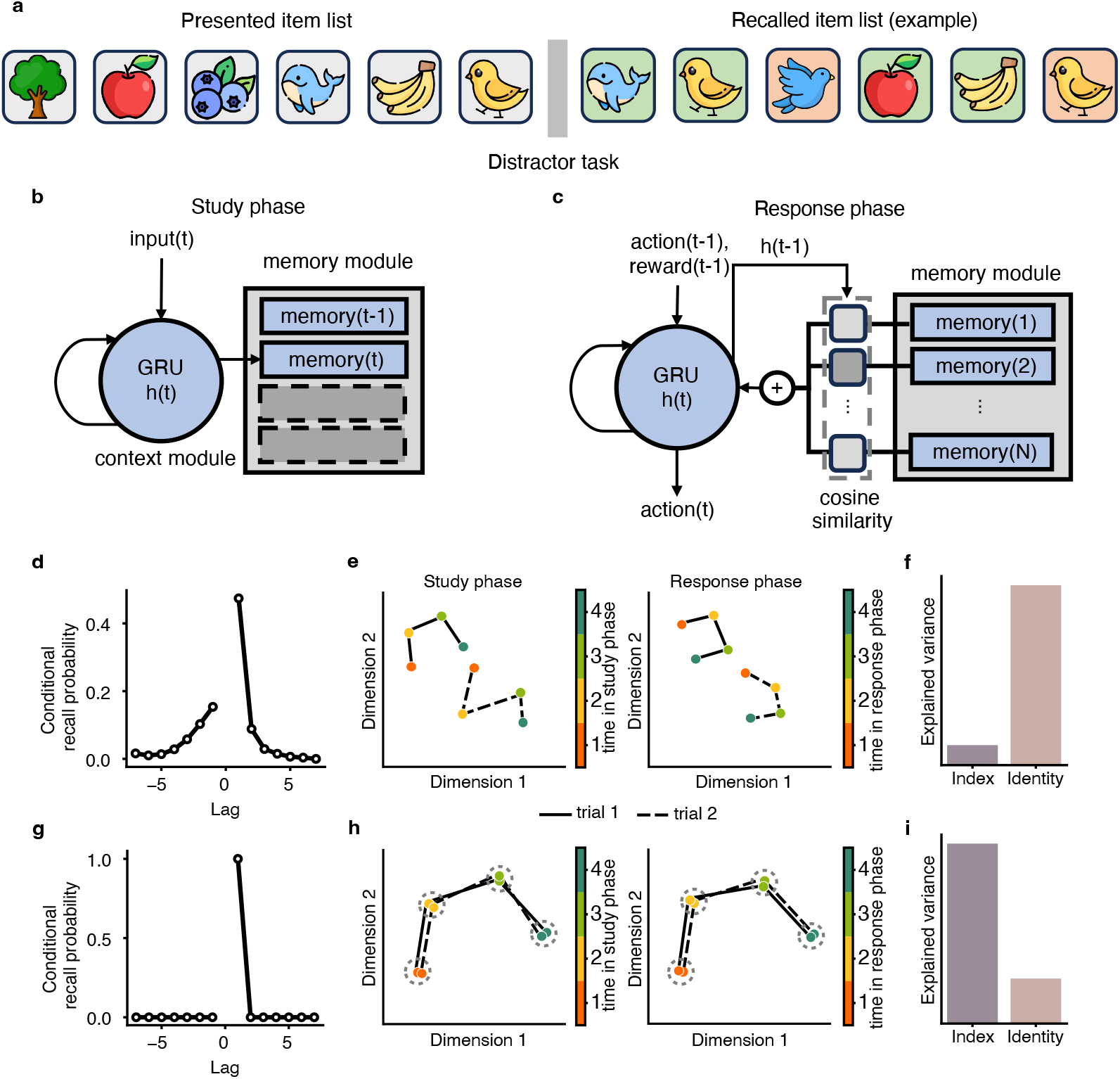
Task and model schematic. (a) Free recall task. The model receives a one-hot encoded item (shown as icons) at each time step during the study phase. In the response phase, it receives a reward for each correct recall (green) and a penalty for each incorrect or repeated recall (red). (b-c) Model architecture. A GRU context module is connected to a slot-based memory module. (b) Study phase: each input item updates the hidden state, which is then appended to the memory module. (c) Response phase: the model computes cosine similarity between the current hidden state and stored memories, retrieving the most similar one. The hidden state is then updated as a function of the previous hidden state, current input, and retrieved memory. Previous action and reward are also provided as inputs during this phase. (d-f) Conceptual illustration of the Temporal Context Model (TCM). (d) TCM predicts a smooth conditional recall probability (CRP) curve: items studied close in time to the last recalled item are more likely to be recalled next, with a forward-order bias. (e) Evolution of the TCM internal state (context) during study (left) and response (right). The state drifts toward each encoded or retrieved item, producing distinct trajectories across trials; recall order can therefore differ from encoding order. (f) Variance in the internal state explained by item index versus item identity in TCM. (g–i) Conceptual illustration of the memory palace technique. (g) The memory palace predicts a high tendency to recall in forward order. (h) Evolution of the internal state in the memory palace. The state represents item index largely independently of item identity, following a fixed trajectory across trials in both study and response phases. (i) Variance in the internal state explained by item index versus identity in the memory palace.

Each trial in the task consisted of a list of 8 items randomly sampled from a dictionary of 64 items, represented as one-hot vectors. During the task’s *study phase*, the model sequentially processes a list of items and stores its internal GRU state in the memory module after each item (Figure 1b). In the subsequent *response phase*, the model uses its current GRU state as a retrieval cue to find the most similar memory in the memory module, recalls the associated item, and updates its state accordingly (Figure 1c). Crucially, to simulate the distractor conditions of human experiments and force reliance on the episodic memory module (Glanzer and Cunitz, 1966), we reset the network’s hidden state between the study and response phases.

To discover effective recall strategies without imposing prior constraints, we trained the model using the advantage actor-critic (A2C) reinforcement learning algorithm (Mnih et al., 2016; Jensen, 2024). The model was rewarded simply for maximizing the number of correctly recalled items and penalized for errors, with no explicit instruction on the order of recall. To determine if any emergent strategies resembled human behavior, we tested whether the model would spontaneously reproduce the temporal contiguity effect—the human tendency to recall items that were studied in close temporal proximity to the just recalled item, and in the forward order (Kahana, 1996). This signature pattern of human recall can be visualized with a conditional recall probability (CRP) curve, which plots the likelihood of recalling an item as a function of its proximity (lag) to the previously recalled item (Figure 1d).

We hypothesized that high recall performance could be achieved through at least two qualitatively different strategies. First, the network might converge on a mechanism based on a drifting temporal context analogous to the Temporal Context Model, using a drifting, item-dependent temporal context to guide recall (Figure 1d-f). This mechanism would be characterized by a smooth CRP curve (Figure 1d), internal states that follow distinct item-dependent trajectories on each trial (Figure 1e), and internal representations dominated by item identity (Figure 1f). Alternatively, the network might discover a more structured, “memory palace”-like strategy that leverages a stable, item-independent scaffold to represent each item’s serial position (Figure 1g-i). This mechanism would lead to a high tendency to recall in the same order as encoded (Figure 1g), stereotyped neural trajectories across trials (Figure 1h), and internal representations dominated by item index (Figure 1i). Distinguishing between these possibilities would therefore reveal the computational principles underlying the model’s emergent solution to the free recall task.

### 2.2 Multiple strategies emerge in a neural network model optimized for free recall

We first asked what computational strategies would emerge when an RNN was given full flexibility to maximize recall performance. After optimizing a diverse population of networks with varying random seeds and hyperparameters, we observed that the networks did not converge on a single solution. Rather, we observed a striking diversity in both behavioral patterns and internal representations. To formally categorize these solutions, we performed *k*-means clustering on the models using metrics that captured both their recall behavior (forward asymmetry, the model’s tendency to recall in a forward order, and temporal organization score, the extent of temporal contiguity) and their internal representations (variance explained by index vs. identity). This analysis revealed the existence of three qualitatively distinct and well-separated strategies for solving the free recall task (Figure 2a). These strategies were distinguished by both their recall behavior—the trade-off between forward asymmetry and temporal contiguity (Figure 2b)—and their internal representations—the balance between coding for an item’s index versus its identity (Figure 2c). Despite their differences, all three strategies achieved high performance (Figure 2d).

**Figure 2:**
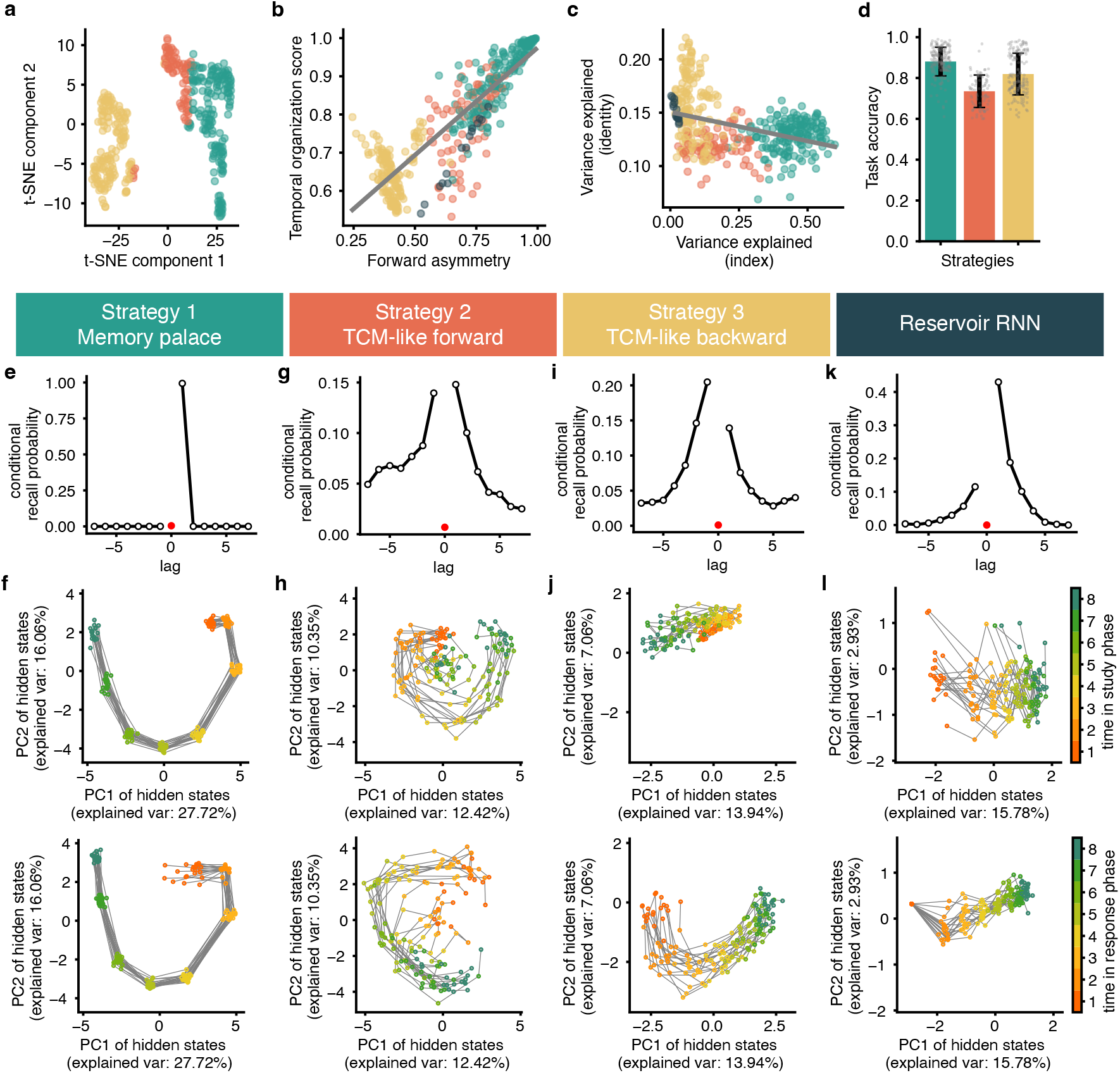
Different strategies of free recall. (a) tSNE embedding of clustering metrics from all trained networks. Different colors denote clusters of models exhibiting strategies 1, 2, and 3 below. (b) Relationship between forward asymmetry and temporal organization score (Pearson correlation, *r*(403) = 0.88, *p <* 0.0001). Black dots indicate reservoir networks, and the other colors indicate trained models exhibiting each of the three strategies. The black line is a linear fit across trained RNNs (excluding the reservoir networks). Forward asymmetry greater than 0.5 reflects a forward-recall bias, while a higher temporal organization score corresponds to a steeper conditional recall probability (CRP) curve. (c) Relationship between hidden state variance explained by item index and hidden state variance explained by item identity (Pearson correlation, *r*(403) = − 0.36, *p <* 0.0001). (d) Mean task performance for each strategy (n=172 for strategy 1, n=78 for strategy 2, n=155 for strategy 3). Error bars denote one standard deviation. (e–l) Representative models for the three emergent strategies and the reservoir RNN. (e, g, i, k) CRP curves showing recall probability as a function of lag relative to the previously recalled item. (f, h, j, l) The first two principal components (PCs) of hidden states during study (top) and response (bottom) phases. Colored dots represent hidden states at each time step; trajectories indicate different trials. A single PCA was fit jointly to study and response phase activity for each model. Each plot shows trajectories from 20 trials with distinct input lists. In the study phase, hidden states are taken immediately after each input. In the response phase, hidden states are taken immediately before memory retrieval (i.e., the state determining which memory is recalled).

To understand the nature of the most dominant high-performance strategy, we analyzed its functional properties. Models employing this strategy exhibited near-perfect, forward-ordered recall, producing a sharp conditional recall probability curve (Figure 2e). Critically, their internal states followed a consistent, stereotyped trajectory during both study and retrieval, regardless of the specific items being processed (Figure 2f). Further analysis showed that an item’s serial index explained substantially more variance in the hidden states than its identity (Figure 3a). Together, these features indicate that the network had learned a stable, item-independent scaffold for memory organization, a mechanism functionally analogous to the “memory palace” technique.

**Figure 3:**
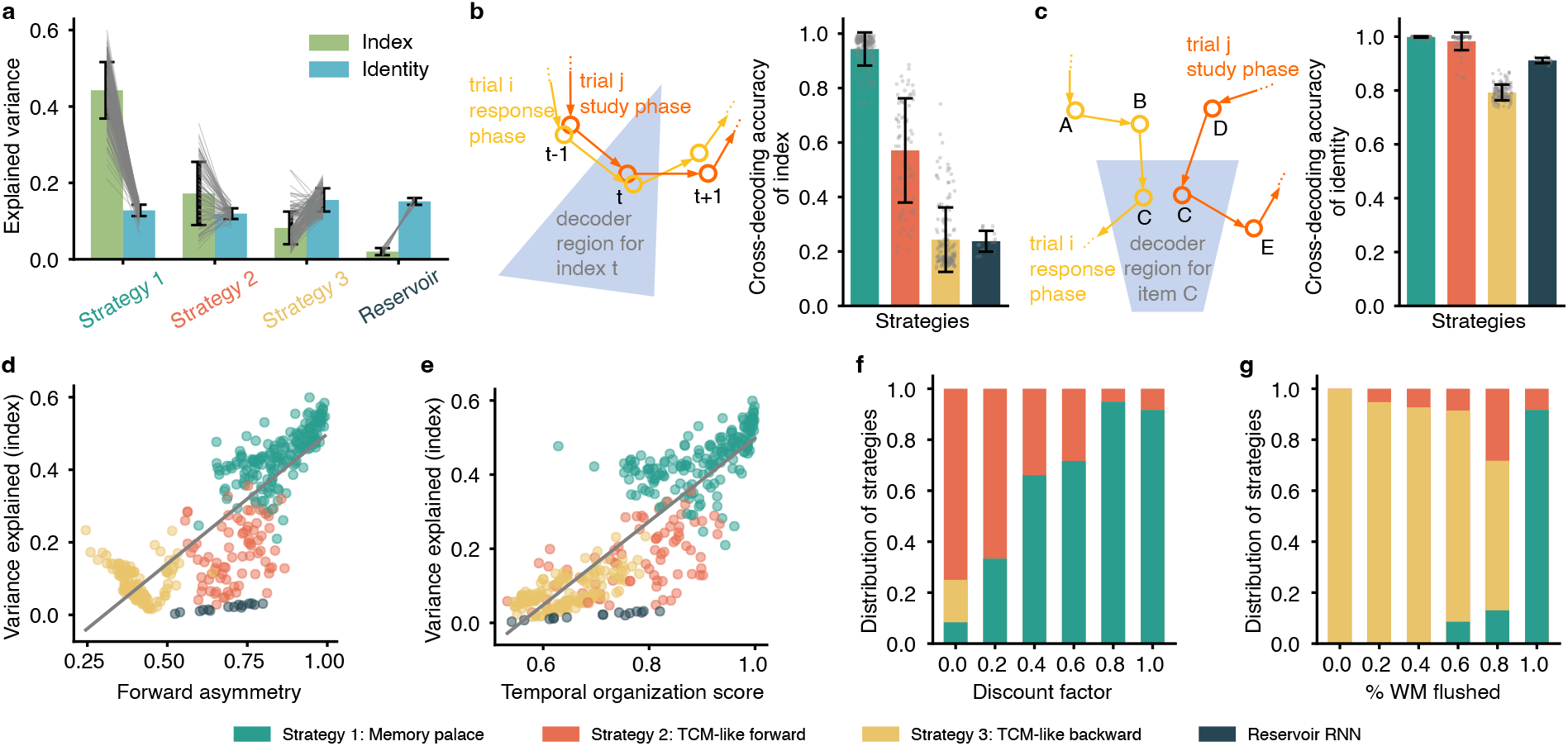
Model representations and factors influencing strategy. (a) Variance in hidden states explained by item index and identity for RNNs with each strategy and the reservoir network. Bars show the average across models, error bars show standard deviation, and lines show individual models (n=172, 78, 155, and 15 for models with strategy 1, 2, 3 and the reservoir network respectively in panels (a)-(c)). (b) Cross-decoding accuracy of item index. A decoder trained on hidden states from study-phase trials with only 8 unique items was tested on response-phase trials with all 64 items, and vice versa. Error bars denote standard deviation and dots denote individual models in panels (b) and (c). (c) Cross-decoding accuracy of item identity. A decoder trained on hidden states from a single study-phase time step was tested across all response-phase time steps, and vice versa. (d-e) Relationship between recall behavior and representational coding. (d) Forward asymmetry versus variance explained by item index (Pearson correlation, *r*(403) = 0.87, *p <* 0.0001). (e) Temporal organization score versus variance explained by item index. Gray lines indicate linear regression fits (Pearson correlation, *r*(403) = 0.87, *p <* 0.0001). (f–g) Training conditions influence strategy distribution. (f) Proportion of models adopting each strategy across different discount factors (20 models per condition; working memory flush held constant). (g) Proportion of models adopting each strategy across different levels of working memory flush (20 models per condition; discount factor held constant).

The memory palace mechanism also emerged when using a more general key-value memory module that supports distinct representations for encoding memory location (keys) and content (values) (Extended data figure 1a,b) (Zhang et al., 2017; Ritter et al., 2018). This generalized architecture can account for some features observed in the human memory system (Gershman et al., 2025; Song et al., 2025). When analyzing the variance in keys and values explained by item index and item identity, we found that keys more strongly represented item index, while values more strongly represented item identity (Extended data figure 1c). This functional separation may play an important role in the formation of the memory palace strategy in the human brain, which could similarly segregate index-based and identity-based representations during structured recall.

We next examined the other two emergent strategies to see how they differed. These strategies closely resembled the mechanisms of the Temporal Context Model (TCM), showing smooth temporal contiguity effects but with opposing directional preferences. The “TCM-like forward” strategy showed a forward recall bias (Figure 2g) and greater trial-to-trial variability in its neural trajectories (Figure 2h). In contrast, the “TCM-like backward” strategy preferred to recall items in reverse order (Figure 2i-j). For both of these strategies, item identity explained more variance than item index (Figure 3a). Thus, our model could not only discover the novel memory palace solution but also learned strategies that operate on principles aligned with existing cognitive models.

To establish a benchmark for these learned solutions, we tested whether recurrent networks would reproduce the temporal context mechanisms of existing cognitive models without learning. We created a “reservoir” network by freezing the random recurrent weights of our model, thereby preventing it from optimizing its dynamics for the task, and trained only its input and output weights. This network developed a strategy that resembled the Temporal Context Model (TCM), exhibiting a classic temporal contiguity effect with forward asymmetry (Figure 2k). An analysis of its internal state revealed that it differed in each trial depending on the specific items presented (Figure 2l), and that it encoded notably more information about item identity than item index (Figure 3a). This result demonstrates that a TCM-like solution is the default behavior for this architecture, confirming that the memory palace is a more specialized strategy discovered through end-to-end optimization.

A critical question was whether the index-based memory palace strategy is fundamentally incompatible with the smoothly drifting dynamics of TCM. To answer that question, we first decoded item identity information from the memory palace network’s hidden states and found that it contained a clear, gradually decaying representation of the currently processed item (Extended data figure 2a,b). Furthermore, an analysis of the hidden states’ cosine similarity revealed smooth temporal dynamics across all strategies, with adjacent states in time being more similar (Extended data figure 2c,d). These findings demonstrate that memory palace and temporal context mechanisms are not mutually exclusive; the network can maintain a stable positional scaffold while simultaneously encoding smoothly drifting content information.

For the memory palace strategy to be effective, its positional code must act as a stable scaffold, remaining invariant across different items (item-invariance) and consistent between encoding and retrieval (phase-invariance). To test this, we evaluated our memory palace models using cross-condition generalization performance (CCGP) (Bernardi et al., 2020). Specifically, we trained a decoder to classify item index from hidden states during the study phase, using trials generated from only 8 random items, and then tested the decoder on hidden states during the response phase with trials containing all 64 items. Models employing the memory palace strategy achieved high cross-decoding accuracy of index, indicating that the same positional code supported both encoding and retrieval across different item sets (Figure 3b). In contrast, reservoir and TCM-like models showed a lower degree of generalization of index representations. This confirms that the memory palace relies on an item-independent index code that can be reinstated during retrieval, analogous to how human experts retrace fixed spatial locations when applying the method of loci. We also assessed the cross-decoding accuracy of identity by training a decoder on hidden states at a single study-phase time step and testing on all time steps in the response phase. Here, all strategies—including memory palace, TCM-like, and reservoir models—showed relatively high cross-decoding accuracy of identity (Figure 3c).

Finally, we sought to determine if a model’s reliance on an index-based code was directly related to its tendency for forward-ordered recall. Across the entire population of trained models, we found a strong positive correlation between the variance explained by index and behavioral metrics of forward recall and temporal contiguity (Figure 3d,e). This result establishes a direct link between the underlying computational mechanism and the resulting recall behavior, showing that models that developed a stronger item-independent scaffold were precisely those that adopted a more systematic, forward-ordered recall policy.

To compare the three RNN strategies with human free recall, we analyzed data from the Penn Electrophysiology of Encoding and Retrieval Study (PEERS), where young adults performed an immediate free recall task (Kahana et al., 2024). By analyzing forward asymmetry and temporal organization scores, we identified a subset of participants who exhibited a strong tendency to recall items in forward order, consistent with the memory palace strategy (Extended data figure 3a). Others displayed smoother temporal contiguity effects with forward asymmetry (Extended data figure 3b), while a third group showed backward asymmetry (Extended data figure 3c). These data suggest that the three strategies identified in our model also reflect naturally occurring recall patterns in humans.

### 2.3 The impact of training conditions on the learned strategies

Having identified multiple distinct strategies in the RNNs trained on free recall, we next sought to understand the conditions under which each strategy emerges. First, we asked how prioritizing long-term rewards over immediate ones would influence the learned strategy. We manipulated this by varying the reinforcement learning discount factor, *γ*, where a high value encourages the model to optimize for recalling the entire list versus than just the next few items. We found that a higher discount factor led to a greater proportion of models learning the memory palace strategy over the TCM-like forward strategy (Figure 3f). This result aligns with findings from human studies, where instructions to recall a whole list encourage participants to initiate recall at the beginning of the list and proceed in a forward order (Tan et al., 2016). Thus, an optimization process that prioritizes long-term, global rewards is a key factor in developing the memory palace strategy.

Second, we asked whether the initial state of working memory at the start of recall would affect the emergent strategy. To simulate the effect of a distractor task—commonly used in human free recall experiments—, we modulated the amount of working memory being “flushed” by injecting a controlled amount of noise into the hidden state between the study and response phases. When working memory was fully preserved (0% flush), the model was biased by the most recent items and consistently adopted the TCM-like backward strategy (Figure 3g). Conversely, when working memory was entirely flushed (100% flush, effectively resetting the hidden state to a random vector), this recency bias was eliminated, and the model overwhelmingly learned the memory palace strategy, initiating recall from the beginning of the list. This suggests that the initial state of working memory is a critical determinant of recall strategy, potentially explaining how humans can improve performance by actively resetting their mental state to overcome the influence of recent events. These results are also consistent with rational analysis of TCM showing that forward-ordered recall is only optimal when retrieval begins from the start of the list (Zhang et al., 2023).

### 2.4 The memory palace strategy is an optimal and robust solution

Having established that our model can learn multiple strategies, we next sought to determine which of them represents the most effective solution for free recall. To ensure a fair comparison, we trained a new set of models with different random seeds under the same discount factor and percentage of working memory flushed and classified their strategies using the cluster centers defined earlier. When comparing performance, the memory palace strategy consistently outperformed both TCM-like forward and backward strategies (Figure 4a). Moreover, task performance was positively correlated with forward asymmetry, temporal organization score, variance explained by index, and cross-decoding accuracy of index (Extended data figure 4a-d) – thereby linking behavioral and representational features of the memory palace to superior recall. Notably, we found a similar trend in the human data. In the PEERS dataset, both forward asymmetry and temporal organization score were positively correlated with free recall performance(Extended data figure 5a,b).

**Figure 4:**
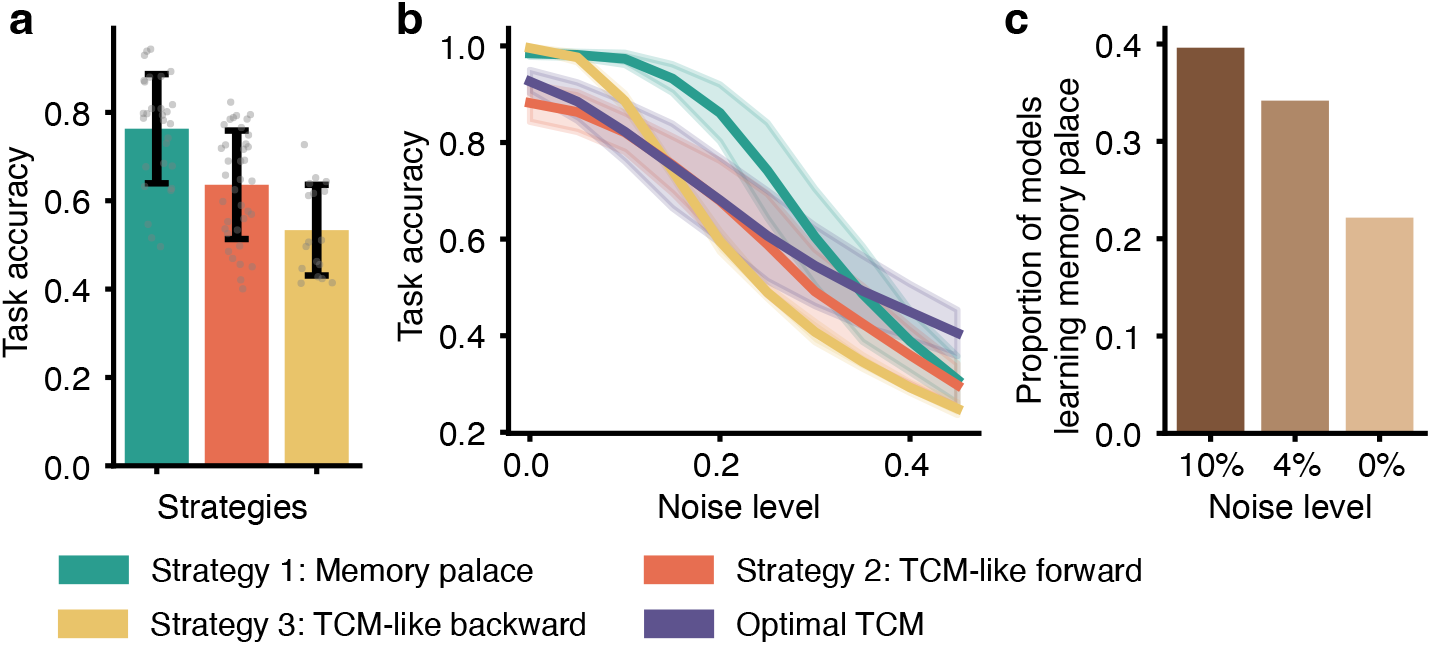
Performance and noise robustness of different strategies. (a) Mean task performance of models within each strategy for a total of 100 models with the same training condition (discount factor = 1.0, proportion of working memory flushed = 0.9). Models were assigned to clusters based on the same cluster centers as in figure 2. Error bars denote the standard deviation. (b) Mean task performance of each strategy as a function of the proportion of noise injected into the recurrent hidden state, together with an optimal TCM model. Error bands denote the standard deviation. (c) Proportion of models that learn a memory palace strategy depending on the proportion of noise injected into the RNN hidden state during training.

We reasoned that a truly optimal strategy must not only be accurate but also robust, so we next asked how resilient each strategy was to neural noise. We tested this by injecting noise into the hidden states of trained models at test time. This analysis revealed that the memory palace strategy was substantially more resilient to noise than either of the TCM-like strategies (Figure 4b). The memory palace strategy outperformed even an optimized TCM model with perfectly forward recall (Zhang et al., 2023). This suggests that the learned index code provides a stable scaffold for memory retrieval that is inherently more robust than a drifting, content-based context, which may be especially important for longer lists.

Given its superior noise resilience, we then asked whether a noisy environment could actively promote the discovery of the memory palace strategy. To test this, we trained new populations of models while adding different levels of noise to their hidden states throughout training. Consistent with our hypothesis, models trained in noisier conditions were more likely to learn the memory palace strategy (Figure 4c). This indicates that the memory palace is not just robust post-hoc, but is preferentially discovered when a network must find a solution that is robust to internal variability.

Taken together, these findings parallel empirical observations in humans, where stronger temporal contiguity and initiating recall from the beginning of a list are associated with better memory performance (Sederberg et al., 2010; Zhang et al., 2023; Romani et al., 2016). The superior performance and robustness of the emergent memory palace strategy therefore support the conclusion that this index-based, forward-recall mechanism represents a computationally optimal solution, providing a normative explanation for its effectiveness in both artificial networks and human memory experts.

### 2.5 External context and stimulus structure shift strategy away from the memory palace

The preceding analyses used one-hot item inputs with no environmental information. However, in more realistic settings, temporally varying external signals—such as background sensory input—can serve as an additional source of temporal context. We therefore asked whether introducing such external information would alter the strategies learned by the model. To test this, we concatenated a slowly drifting Gaussian bump vector to the item input as external context (Figure 5a). This signal was independent of item identity and drifted by one dimension per time step. We trained models across a range of context amplitudes and found that stronger external context progressively shifted models away from the memory palace strategy: conditional recall probability curves became flatter (Figure 5b), more models adopted TCM-like strategies (Figure 5c), and temporal organization scores, forward asymmetry, and index representations all decreased (Figure 5d-g). These results indicate that when external temporal context is available as an input, models rely on it rather than learning an internal positional scaffold.

**Figure 5:**
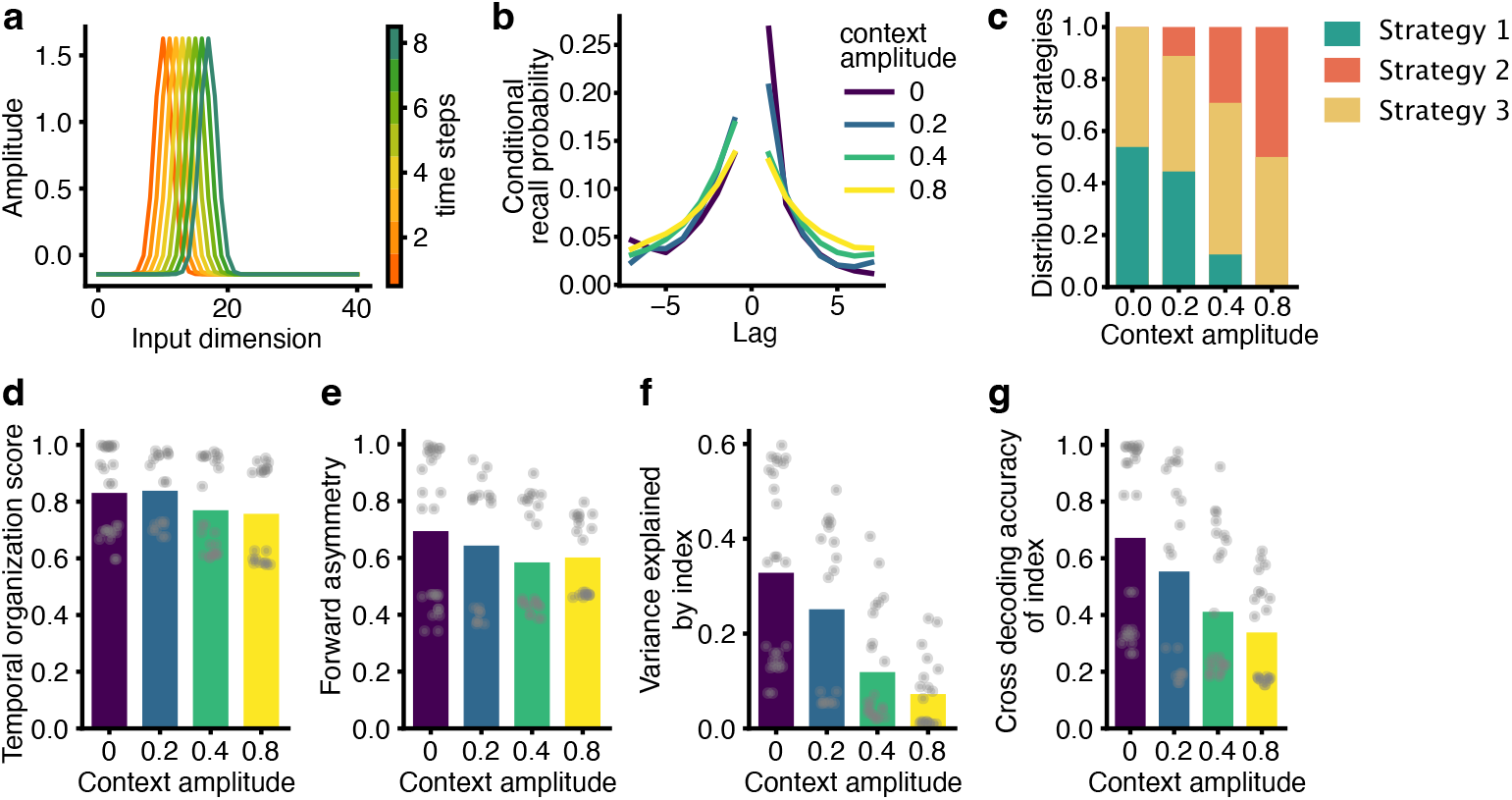
External temporal context shifts models toward TCM-like strategies. All panels show results from 20 models per condition, trained with the same settings as Figure 4 (discount factor = 1.0, proportion of working memory flushed = 0.9). Strategies were classified using the same features and cluster centers as in previous figures. (a) Example sequence of external context vectors during the study phase. The context is a 40-dimensional Gaussian bump that shifts by one dimension per time step, producing a slowly drifting temporal signal. The starting position is randomized on each trial. (b) Mean conditional recall probability curves for models trained with different context amplitudes (0, 0.2, 0.4, 0.8). (c) Proportion of models adopting each strategy as a function of context amplitude. (d) Mean temporal organization score, (e) forward asymmetry, (f) variance in hidden states explained by item index, and (g) cross-decoding accuracy of index from hidden states, each as a function of context amplitude. Dots represent individual models (n=20 for each context amplitude).

While external signals can guide the model to rely on temporal context, it may potentially encourage memory palace by providing a stable location-like signal. We hypothesize that this would happen when matching context is provided across encoding and retrieval. To test this, we trained models with either matching or mismatched context across the study and response phases (Extended data figure 6a,b). Models trained with matching context performed better and were more likely to adopt the memory palace strategy than those trained with mismatched or no context (Extended data figure 6e-i). Switching from matching to mismatched context at test time reduced both performance and memory palace adoption, whereas the reverse switch—training with mismatched context but testing with matching—produced no apparent benefit. Moreover, fixing the same context sequence across trials and phases further increased memory palace adoption. Together, these results suggest that consistent external context across encoding and retrieval can function as location information, guiding the model to learn the loci of the memory palace and subsequently cueing recall. This is consistent with established findings that encoding and retrieval in the same environment improves memory performance (Smith and Vela, 2001).

Beyond temporal context, naturalistic stimuli also carry semantic structure that could influence retrieval strategy. We therefore trained models with a two-level hierarchical semantic embedding for each item, in which items shared features at two scales of granularity (Extended data figure 6c,d). As the amplitude of semantic information increased, models showed reduced temporal contiguity (Extended data figure 6j) and greater semantic contiguity, recalling semantically similar items in succession (Extended data figure 6k). Models also became less likely to adopt the memory palace strategy (Extended data figure 6l-n). Thus, when items carry shared semantic features, models exploit this structure for retrieval rather than relying on a purely positional code.

### 2.6 The optimal strategy depends on the task demands

Although memory palace appears to be a highly effective strategy for free recall in both our simulations and human behavior, its scope of optimality remains unclear. We hypothesized that this strategy is particularly well suited to free recall, where success depends only on retrieving labels rather than content. By contrast, everyday memory often requires flexible retrieval based on an item’s features or identity, using the content within each memory for downstream tasks. In such settings, we predicted that a purely index-based scaffold would be ill-suited.

To test this hypothesis, we trained models on a “conditional free recall” task variant that required content-based retrieval. In this task, each item was assigned a set of features, and during the response phase the model was cued to recall only items with a specific feature value (Figure 6a). We trained the model under the same conditions that strongly promoted the memory palace strategy in the free recall task (i.e., a high discount factor and full working memory flush). We then tested this model on a standard free recall task to analyze the learned strategy. We found that these models achieved moderate performance on both the conditional and standard free recall tasks (Figure 6b). Notably, they exhibited all the hallmarks of a TCM-like strategy: a smooth temporal contiguity effect (Figure 6c), variable trial-to-trial neural trajectories (Figure 6d), and internal representations dominated by item identity rather than index (Figure 6e). None of the models learned the memory palace strategy. These results demonstrate that the optimality of the memory palace strategy is task-dependent, representing a highly specialized solution for sequential recall that is discarded when flexible, content-based retrieval is required.

**Figure 6:**
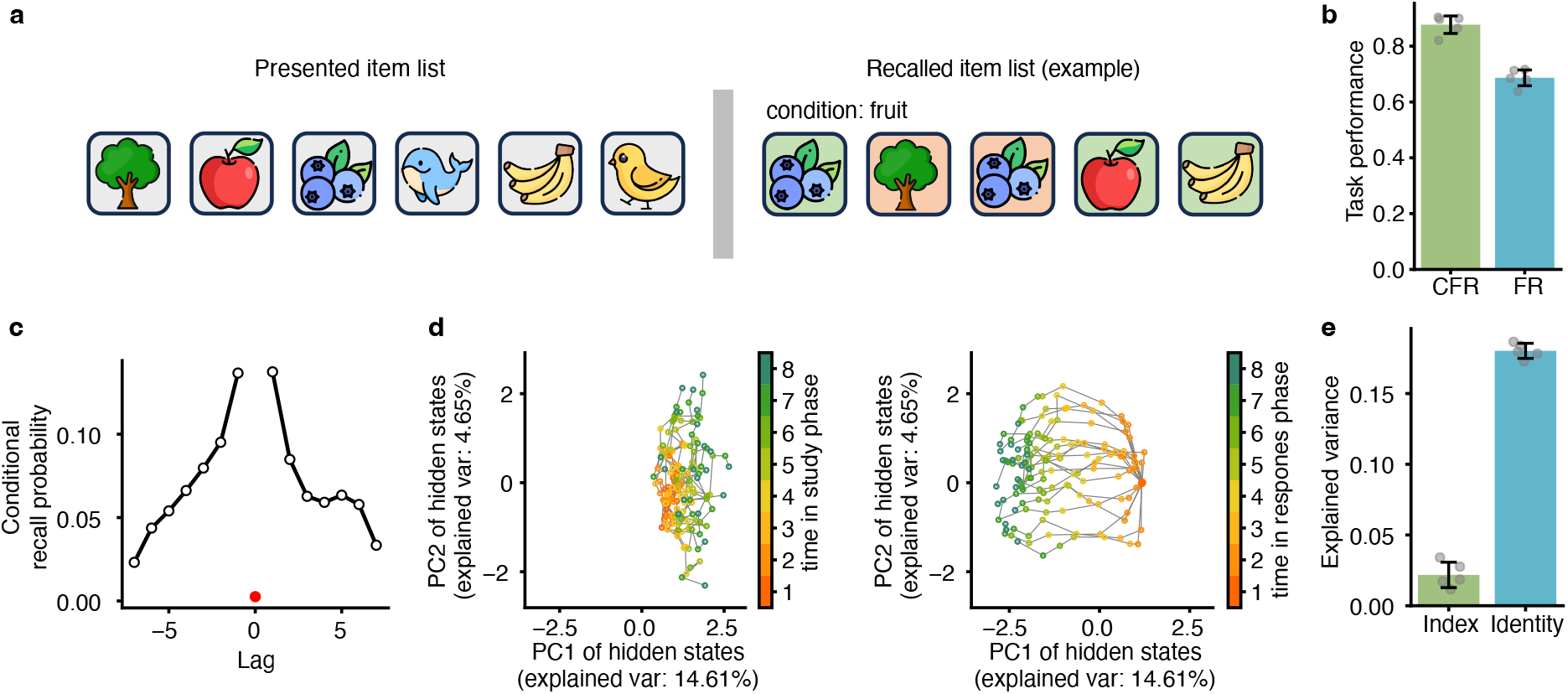
Content-based memory search discourages the memory palace strategy. (a) Schematic of the conditional free recall task. Each item was assigned a set of features, and during the response phase the model was cued to recall only items with a specified feature value. Correct recalls (green) were rewarded, while errors (red) occurred if the recalled item was absent from the list, repeated, or failed to match the condition. (b) Task performance of models trained on the conditional recall task and subsequently tested on standard free recall. Bars show the mean across 5 models trained with different random seeds; error bars denote one standard deviation. (c) Conditional recall probability (CRP) curves showing recall likelihood as a function of lag from the previously recalled item. (d) The first two principal components (PCs) of GRU hidden states during the study phase (left) and response phase (right). Each trajectory represents a trial. (e) Fraction of variance in hidden states explained by item index versus item identity. Error bars denote one standard deviation and dots show each individual model (n=5).

## 3 Discussion

This study demonstrates that a neural network optimized for free recall spontaneously discovers multiple, distinct strategies for solving the task. While some of these strategies were reminiscent of the drifting temporal context assumed by standard cognitive models (Howard and Kahana, 2002; Sederberg et al., 2008), others developed an alternative mechanism: an item-independent scaffold that encodes each item’s sequential index to guide systematic forward retrieval. This “memory palace” strategy yielded superior performance and robustness, and its emergence was favored by conditions that prioritized whole-list recall and discouraged reliance on working memory. These results show that human-like recall patterns can arise from different computational principles and suggest that the memory palace represents a normative, computationally optimal solution for free recall.

Our modeling approach provides a normative explanation for the effectiveness of systematic recall strategies, extending previous rational analysis of memory (Zhang et al., 2023). By training a network without built-in biases, we showed that a memory palace strategy emerges not as a contrived mnemonic, but as a high-performing, optimal solution. This finding offers a mechanistic basis for the empirically observed link between strong temporal contiguity and better recall performance in humans (Sederberg et al., 2010; Romani et al., 2016), supporting theoretical claims that forward-ordered recall is an optimal policy (Zhang et al., 2023). Our representational analyses further connect this behavior to the formation of an item-invariant index code, akin to the mental locations used by memory experts. Therefore, our results suggest that the memory palace is not merely a contrived mnemonic but may be a general and computationally optimal strategy for free recall.

The factors that promoted the memory palace strategy in our model may also offer testable hypotheses about the origins of individual differences in human recall. We found that a higher reinforcement learning discount factor—the tendency to prioritize future over immediate rewards—was a key determinant of this strategy’s emergence. This suggests that individuals more disposed to long-term planning may be more likely to develop systematic, forward-ordered recall strategies, a hypothesis testable in future experiments (Basile and Toplak, 2015; Sadeghiyeh et al., 2020). Similarly, the fact that the index-based strategy was learned most effectively when recency effects were suppressed suggests that the ability to overcome the influence of immediately preceding events is a critical component of developing an optimal recall policy.

A key conclusion of our study is that the optimality of a memory strategy is highly task-dependent. While our work highlights the memory palace as a powerful solution for free recall, its dominance vanished when task demands shifted to require flexible, content-based search. As demonstrated in our external context and conditional free recall experiments, models also shifted towards TCM-like strategies when stimuli contained a strong external context or when retrieval had to be guided by item features rather than sequential position. This suggests that the memory palace is a useful but specialized strategy for retrieval of independent items, while TCM-like dynamics may better support flexible adaptation to diverse problems (Bennion et al., 2025). Furthermore, our model clears its memory across trials, which does not account for situations that require multiple separate sequences to be kept in memory. Humans distinguish between different events using event context. This could make the memory palace less optimal, since items stored at the same location may interfere with one another in the absence of an event context. Future work could test how strategy choice shifts in tasks requiring integration or contextualization of many items, which may be more useful for guiding future actions than precise episodic detail.

The index code learned by our model can be understood as an abstract form of loci: each serial position in the list serves as a distinct, stable location onto which items are bound during encoding and systematically retraced during retrieval. In practice, humans using a memory palace technique often rely on a richer two-dimensional spatial scaffold grounded in personally meaningful environments, where perceptual detail and vivid imagery further strengthen item-location associations. A natural extension of our work would be to integrate our framework with architectures that learn relational memory by analogy to spatial map formation (Whittington et al., 2020; Chandra et al., 2025). This could yield a spatially-grounded architecture that preserves the temporal retrieval dynamics while also supporting the richer associative structure that characterizes expert-level human memory performance.

Finally, our model provides a powerful and flexible framework for exploring how episodic memory supports broader cognitive functions. Because the model learns human-like retrieval patterns from basic principles— without hand-crafted cognitive components—it can be readily adapted to study how memory supports higher-order cognition like decision-making, planning, and inference. For instance, future work could extend this model to planning tasks that require retrieving sequences of memories to simulate future outcomes. Such research could bridge our understanding of memory retrieval and its ultimate function in complex, goal-directed behavior.

## 4 Methods

### 4.1 Task

#### Free recall

We used a free-recall task to train the model. There was a study phase and a response phase in each trial. During the study phase, the model was presented with a list of items. During the response phase, the model was trained to recall the studied items in any order. Items were represented as 64-dimensional one-hot vectors, and each trial contained a list of 8 items as the input sequence.

More specifically, during the study phase, the model received one item per time step as input. During the response phase, the model recalled an item per time step as output. A correct output resulted in an immediate reward (+1) to the model. If the model recalled an item that was not presented during the study phase, or if it recalled an item more than once, the outputs were marked as incorrect and the model received a penalty (-0.75). The item list presented during the study phase was generated independently on each trial by randomly sampling 8 items from the full set of 64 items without replacement. There was a time limit of 8 time steps in the response phase, and the model needed to recall all the items correctly to receive the full reward.

#### Conditional free recall

In the conditional free recall, 6 features were assigned to each item. Each feature had 2 values, which was represented as a one-hot vector. Therefore, there are still 64 items in total. During the study phase, there was 8 items presented as 64-dimension one-hot vectors for each trial. During the response phase, we gave the model a condition as input. The condition was in the form of “feature x equals to y”, in which x takes the value of 1 to 6, and y takes the value of 1 or 2, both encoded as one-hot vectors. The models needed to recall only items in the list that match this condition. A correct recall led to a reward of +1 and a wrong or repeated recall led to a penalty of -0.75. The output was a 64-dimensional vector representing the probability distribution of recalling each item, same as the free recall task. The model had 8 time steps to recall for each trial. It could take an action representing “no action” for not outputting any items, and it got no reward or penalty for it. To avoid the model from always taking no action, the model got a penalty of -0.75 for each not-recalled item at the end of the trial if there were items that matched the condition but were not recalled.

The model was trained on this conditional free recall task. During testing, we tested the model on a normal free recall task by setting a feature in all items of the list to be the same and setting the condition to be this feature and its corresponding value.

### 4.2 Model structure

The model was composed of a ‘context module’ and a ‘memory module’. The ‘context module’ consisted of 128 gated recurrent units (GRU) (Cho et al., 2014). At each time step, the GRU received inputs and produced outputs with evolving dynamics according to

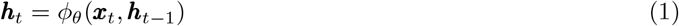

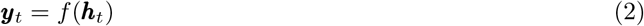

Here, *ϕ* denotes a forward pass through the recurrent GRU dynamics, *θ* denotes the set of all model parameters, ***x***_*t*_ are inputs to the GRU, and ***y***_*t*_ are outputs computed from the GRU hidden state ***h***_*t*_. The ‘memory module’ consisted of a list of slots for storing memories. At the start of each trial, the memory module was cleared to allow the storage of new memories, and the hidden state was reset to all zeros.

For each time step in the study phase, the GRU received the one-hot representation of a study item. It then updated the GRU hidden state ***h***_*t*_ and appended it to the memory module as an episodic memory ***m***_*t*_.

In the response phase, the model retrieved a memory from the memory module and produced an output at every time step. After retrieving a memory, the GRU took the retrieved memory as part of its inputs and updated its hidden state based on the retrieved memory as well as other inputs and the previous hidden state. The output consisted of a 64-dimensional policy *π*_*θ*_(*a*_*t*_), which was a set of probabilities associated with each possible item to recall. An action *a*_*t*_ was sampled from the policy as the recalled item.

The memory retrieval process was as follows. At the start of each time step, the model computed the cosine similarity between the current hidden state ***h***_*t−*1_ and the *N* memories stored in the memory module {***m***_*i*_|*i* = 1, …, *N* } to produce a similarity vector ***s***_*t*_,

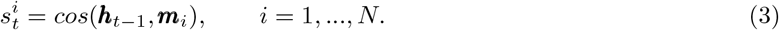

The model then computed the retrieved memory as the weighted sum of all the stored memories according to the weights of the similarity vector. To favor the retrieval of a single memory rather than a combination of multiple memories, we first passed the similarity vector through a softmax function with a low temperature *τ*,

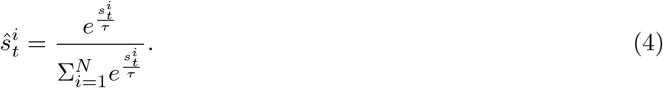

Then we computed the retrieved memory 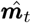 by treating 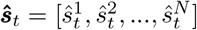 as weights for each memory,

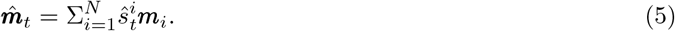

This weighted averaged memory was used instead of the single most similar memory during training to make the memory retrieval process differentiable during training. The softmax temperature of computing memory similarity *τ* gradually decreases from 0.01 with a rate of 0.9 every 32,000 episodes until reaching less than 0.0002. During testing, we made the model retrieve the single most similar memory instead of a weighted sum to prevent the model from retrieving a uniform combination of all memories. Through gradually reducing the softmax temperature during training, the performance was not affected during testing.

After retrieving a memory, the GRU updated its hidden state on the basis of the retrieved memory as well as the input and previous hidden state.

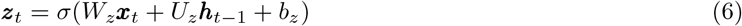

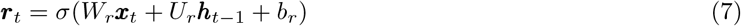

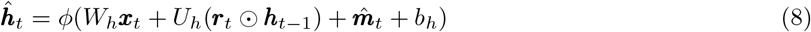

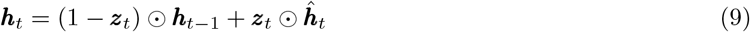

To force the model to use previously stored episodic memories in the memory module instead of working memory stored in the hidden state, the hidden state of the GRU was reset to random noise between the study and response phases by default (proportion of working memory flushed equals to 1). We allowed the model to update its hidden state for a single time step without any recall before retrieving the first memory, so that it could learn an initial state for retrieving the first memory.

### 4.3 Training algorithm

We trained the model on the free recall task with the advantage actor-critic (A2C) reinforcement learning algorithm (Mnih et al., 2016; Jensen, 2024). The parameters *θ* updated in the form

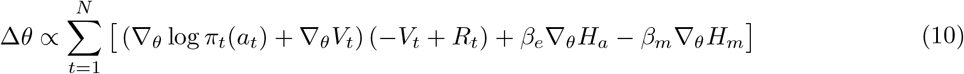

Here, *π*_*t*_(*a*_*t*_) is the policy output by the model at time step *t, a*_*t*_ is the randomly sampled action from the policy (i.e. the item recalled), *V*_*t*_ is the value estimated by the network, and 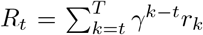, where *r*_*t*_ is the reward at time step *t* and *γ* is the discount factor. *H*_*a*_ = −∑ _*a*_ *π*_*t*_(*a*_*t*_) log *π*_*t*_(*a*_*t*_) is the entropy of the policy, and *β*_*e*_ is the weight of the entropy regularization term.

The last reward *r*_*t−*1_ and last action *a*_*t−*1_ sampled from the policy *π*_*t−*1_(*a*_*t−*1_) were returned to the model as inputs at the next time step (Wang et al., 2018). This is important for the model to perform the task, since it provides information about what items have been previously recalled, and therefore also which items have yet to be recalled.

To encourage the model to recall only one memory at a time and to inhibit the retrieval of other memories, we added another entropy regularization term 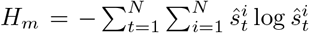on the recall weights **ŝ**_*t*_ in the loss function. This term encouraged the model to decrease the entropy of the memory similarity **ŝ**_*t*_, which made it closer to a one-hot vector. As a consequence, the retrieved memory 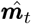 will be closer to a single memory in the memory module instead of a combination of multiple memories.

We trained the neural network model using the Adam optimizer for 1.6 × 10^6^ episodes with a batch size of 16 and a learning rate of 0.003.

### 4.4 Varying training settings

When we investigated the influence of the proportion of noise injected into working memory between phases, the initial hidden state at the start of the response phase was computed as

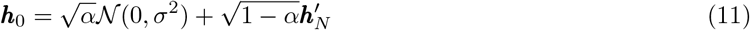

Here 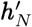 is the last hidden state during the study phase. N (0, *σ*^2^) is sampled from a normal distribution in which *σ* is the standard deviation of 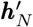. This computation maintains the variance of the hidden state, and *α* is the proportion of variance replaced by noise in the hidden state, varying from 0 to 1.

We varied the discount factor *γ* and the noise proportion *α* from 0 to 1 with an interval of 0.2. For each combination of hyperparameters, we trained 20 models with different random seeds.

When adding noise to the working memory to analyze the noise robustness of the models (Figure 4b,c), we used the same method as above. When noise was added to trained models for testing (Figure 4b), we varied noise proportion from 0 to 0.5 with an interval of 0.02. When noise was added to models during training (Figure 4c), we chose noise proportions of 0.1, 0.04 and 0. The same group of models with different discount factors and noise between phases was trained with different proportions of working memory noise.

### 4.5 Data analyses

#### 4.5.1 Behavioral metrics for contiguity effect

##### Conditional recall probability

The conditional recall probability (CRP) curve is a basic method for illustrating the contiguity effect. For an item *x*_*i*_ at position *i* in the presented list, we computed the probability of recalling the item *x*_*i*+*j*_ immediately after recalling the item *x*_*i*_, in which *j* is the lag of item *x*_*i*+*j*_ to the item *x*_*i*_ in the original list. This way we got a probability distribution of recalling the next item as a function of *j*. We then summed up this distribution for all items by aligning at lag 0, and divided it by a random recall distribution, so that we got the CRP curve.

##### Forward asymmetry

We used the forward asymmetry and the temporal organization score to evaluate the contiguity effect of a model. The forward asymmetry (FA) is the tendency to recall in the forward order compared to the backward order. It is defined as the proportion of forward recall transitions in all recall transitions, where a recall transition is defined as a pair of consecutive recall instances. We define RT(*i, j*) as a recall transition corresponding to first recalling the *i*th item and then the *j*th item in the studied list, and the forward asymmetry can then be computed as

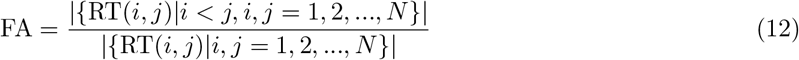

A forward asymmetry of 1 means perfect forward recall, 0 means perfect backward recall, and 0.5 means no bias in recalling forwards or backwards.

##### Temporal organization score

The temporal organization score (TS, also named as temporal factor) was introduced by Polyn et al. (2009a), and is a widely used metric that quantifies the tendency of an agent to recall an item that is in close temporal proximity to the previously recalled item (Sederberg et al., 2010; Hong et al., 2024). The temporal organization score of a recall transition RT(*i, j*) in a given sequence of recall behavior is defined as

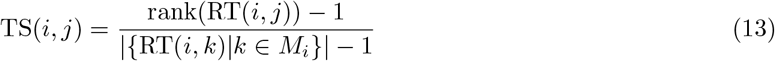

Here *M*_*i*_ = [1, *N* ] ∩ ℤ\{*m*_*t*_|*t* = 1, 2, …, *i*} is the set of the positions of all items that have not yet been recalled in the list. Therefore, {RT(*i, k*)|*k* ∈ *M*_*i*_} is the set of all possible correct transitions after recalling the *i*th item. rank(RT(*i, j*)) is the rank of RT(*i, j*) in the set RT(*i, k*) |*k* ∈ *M*_*i*_ *}*, ordered according to the negative absolute value of serial position lag (− |*j*− |*i* ). When there is a tie, the rank of all members of the tie will be the mean of their ranks. The closer *i* and *j* is, the higher TS(*i, j*) will be. The temporal organization score of a model is the mean score across all recall transitions. Invalid recall transitions were not accounted, such as transitions with recalling a false item or a repeated item. A temporal organization score of 1 refers to perfect contiguity, i.e. the model always recalls the adjacent item to the just recalled item in the presented list. A temporal organization score of 0.5 refers to no temporal contiguity, with a uniform probability of recalling items in any order.

#### 4.5.2 Representational analyses of strategy

To quantify the strategy of a model, we used both dimensionality reduction methods and decoding methods. We recorded 5,000 free recall trials from each trained model to perform the analyses.

##### PCA

We did PCA on the hidden state of the model by concatenating the hidden state in the study phase and the response phase. We fit the PCA on the 5,000 trials and plotted 20 trials for each model.

##### Decoding index and identity

We used Ridge classifiers to decode the item identity and item index information from the hidden states at each time step. We trained different classifiers for decoding the item identity of items at each time step from GRU hidden states at each time step. For the study phase, items that were studied at a certain time step were combined across trials. For the response phase, items that were recalled at a certain time step were combined across trials.

##### Explained variance of index and identity

We used linear regression to fit the hidden state with the corresponding index and identity.

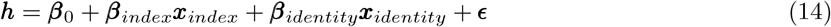

in which ***h*** is the hidden state, ***β***_index_ and ***β***_identity_ are the weights for index and identity, ***β***_0_ is the bias, and ***ϵ*** is the residual error. ***x***_index_ and ***x***_identity_ are one-hot representations of index and identity. We then quantified the variance of hidden state explained by index and identity.

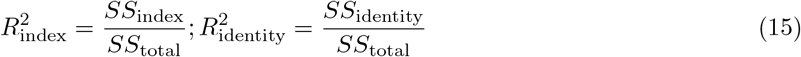

in which *SS*_total_ is the total sum of squares of the hidden state samples recorded during testing the model, *SS*_index_ and *SS*_identity_ are the regression sum of squares with index and identity. As the index and the identity are independent of each other, the explained variance of the two variables does not overlap. The regression was performed separately for the study phase and response phase, and we averaged the explained variance across the two phases. For the study phase, the identity for each hidden state was the item currently presented, and the index was the presented order of the item. For the response phase, the identity was the currently recalled item, and the index was the index of the recalled item in the original list.

##### Cross-decoding index and identity

For models using the memory palace strategy, hidden representations of index for two different items (from two different trials) with the same index should be similar, and hidden representations of index for the study phase and the response phase should be similar. We measured the level of this item-invariance and phase-invariance of the index code through decoding (Bernardi et al., 2020). For each model, we trained an SVM decoder on hidden states in the study phase with a set of trials including only 8 unique items from the total of 64 items. Then the decoder was tested on hidden states in the response phase with trials generated with all items. We trained another decoder on the response phase and tested on the study phase, and averaged these two decoding results to get the final cross-decoding accuracy of index. We also examined the cross-decoding accuracy of identity. The decoder was trained on hidden states in the study phase with only one same time step from each trial, and was tested on hidden states in the response phase with all time steps of each trial, and vice versa.

#### 4.5.3 Clustering for strategies

To discover the strategy from the space of the models, we performed a k-means clustering based on the following 9 metrics of each model: forward asymmetry, temporal organization score, decoding accuracy of current index, decoding accuracy of current identity, decoding accuracy of the last item identity, explained variance by index and identity, and cross-decoding accuracy of index and identity. Clustering was performed on all models with varying discount factor, proportion of working memory flushed between phases and noise injected to the hidden state during training. We filtered out models with free recall performance less than 60%. We tried different numbers of clusters, and computed the inertia and the Sihouette score of each number of clusters to identify the proper number of cluster (see Supplementary figure 3). We chose a cluster number of 3 based on the elbow method, which identifies the point where the decrease in inertia began to slow, and referred to the Sihouette score.

### 4.6 The key-value memory model

We trained a model with learnable key/value representations to investigate if our results can be extended to commonly-used key-value memory in machine learning. During the study phase in the key-value memory model, the hidden state passed a feedforward layer to form a key, and passed another feedforward layer to form a value. Each key-value pair was a memory (Extended data figure 1a). During the response phase, the hidden state passed the same feedforward layer as the key to form a query, and retrieved the value with the highest cosine similarity between the query and the corresponding key. We found that using the same feedforward layer to generate keys and queries is important for contiguity effect and the memory palace strategy to emerge from the model. If the feedforward layer is different for keys and queries, the model cannot show contiguity effect in the first place.

### 4.7 Human data

We used data from the Penn Electrophysiology of Encoding and Retrieval Study (PEERS) (Kahana et al., 2024). The behavioral analysis included 200 young adult participants aged between 18 and 30 who have completed experiment 1 in the dataset, which was an immediate free recall task. We extracted trials with 16 items in a word list, and split trials for each participant to two halves for cross-validation. We computed forward asymmetry and temporal organization score for each participant with the first half of trials using the same algorithm described above. Based on observation in model clusters, we matched participants with a temporal organization score larger than 0.8 to the memory palace strategy, and then split participants to TCM-like forward and TCM-like backward strategies by comparing their forward asymmetry with 0.5. We randomly selected 4 participants from each group as examples that match to each strategy found in the model, and plotted the conditional recall probability curve with the second half of the trials for each participant in Extended data figure 3.

### 4.8 External context and semantic stimuli

The external temporal context stimulus was a 40-dimensional Gaussian curve, shaped as the PDF of a normal distribution over the range [-3, 3] with mean 0 and standard deviation 0.2. When used as model input, it was concatenated to the one-hot item embeddings and scaled by an amplitude between 0 and 0.8. To simulate gradual contextual drift over time, the peak of the Gaussian curve shifted by one dimension per time step. This context sequence for each trial was generated randomly.

By default, the context vector was included as input only during the study phase (for Figure 5). We then simulated cases where both the study phase and the response phase have external context inputs. When context was matched across phases, the identical sequence of context vectors was presented in both the study and response phases. When context differed across phases, a new sequence was generated for the response phase, with peak dimensions that diverged from those of the study phase (Extended data figure 6a).

Semantic information was represented as a 20-dimensional vector with a one-to-one mapping to each item, comprising two features. The first feature was 4-dimensional and shared across every 16 items; the second was 16-dimensional and shared across every 4 items (Extended data figure 6c). Gaussian noise (*σ* = 0.1) was added to each vector. This produced a two-level hierarchical semantic structure across all items, with a similarity matrix shown in Extended data figure 6d. When presented as inputs to the model, semantic vector was concatenated to the original one-hot embedding of items and was scaled by an amplitude of 0 to 0.5.

### 4.9 Data availability

The Penn Electrophysiology of Encoding and Retrieval Study (PEERS) dataset (Kahana et al., 2024) used for comparison with human data is publicly available at https://memory.psych.upenn.edu/files/PEERS.data.tgz.

### 4.10 Code availability

Code for this paper is available via GitHub at https://github.com/Veritaria/rnn-free-recall (Li, 2026).

## 5 Extended data

**Extended Data Figure 1:**
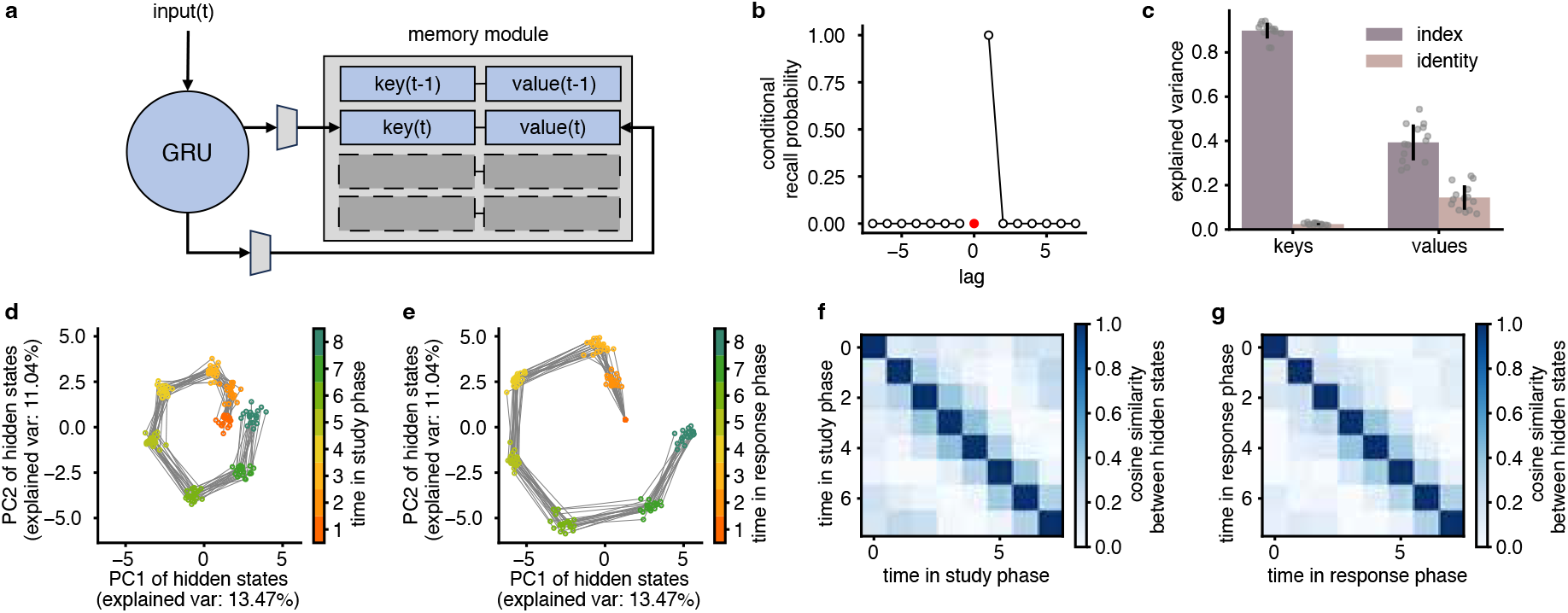
A key-value memory network showed the memory palace strategy. (a) An illustration of the key-value memory model structure. During the study phase, the GRU hidden state was converted to a key and a value through two different feedforward layers. During the response phase, the GRU passed its hidden state to the same feedforward layer as generating the keys to form a query vector, computed the cosine similarity between the query and all keys, and retrieved the value with the maximum similarity. (b) The conditional recall probability of the key-value memory model with default training settings the same as the previous RNN-memory model. (c) Explained variance of index and identity in keys and values. The keys contained more index information than the values, while the values contained more identity information than the keys. 20 models with different random seeds were included (n=20). Error bars denote one standard deviation and dots denote individual models. (d) PCA trajectories of the hidden state in the study phase. (e) PCA trajectories of the hidden state in the response phase. (f) Cosine similarity between each key. (g) Cosine similarity between each value.

**Extended Data Figure 2:**
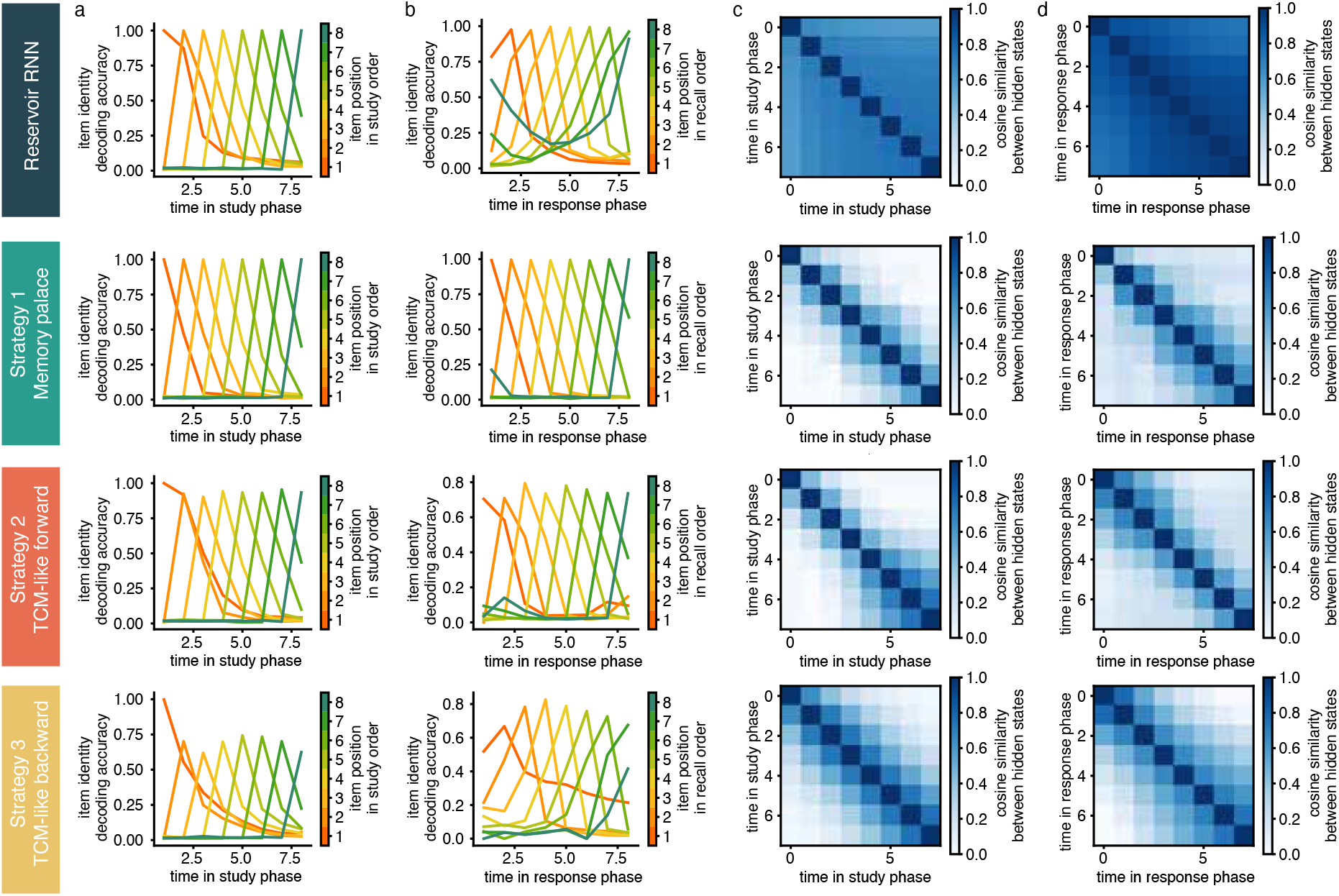
Additional analysis for each strategy. (a) Decoding accuracy of item identity from hidden states during the study phase. Each curve represents items presented at a particular time step. Each data point in the plot is generated with a different decoder, which is trained with cross-validation on data from 5000 trials. (b) Decoding accuracy of item identity from hidden states during the response phase. State similarity between each time step in the study phase. (d) State similarity between each time step in the response phase. For all strategies, the hidden state changes smoothly during both study and response phases.

**Extended Data Figure 3:**
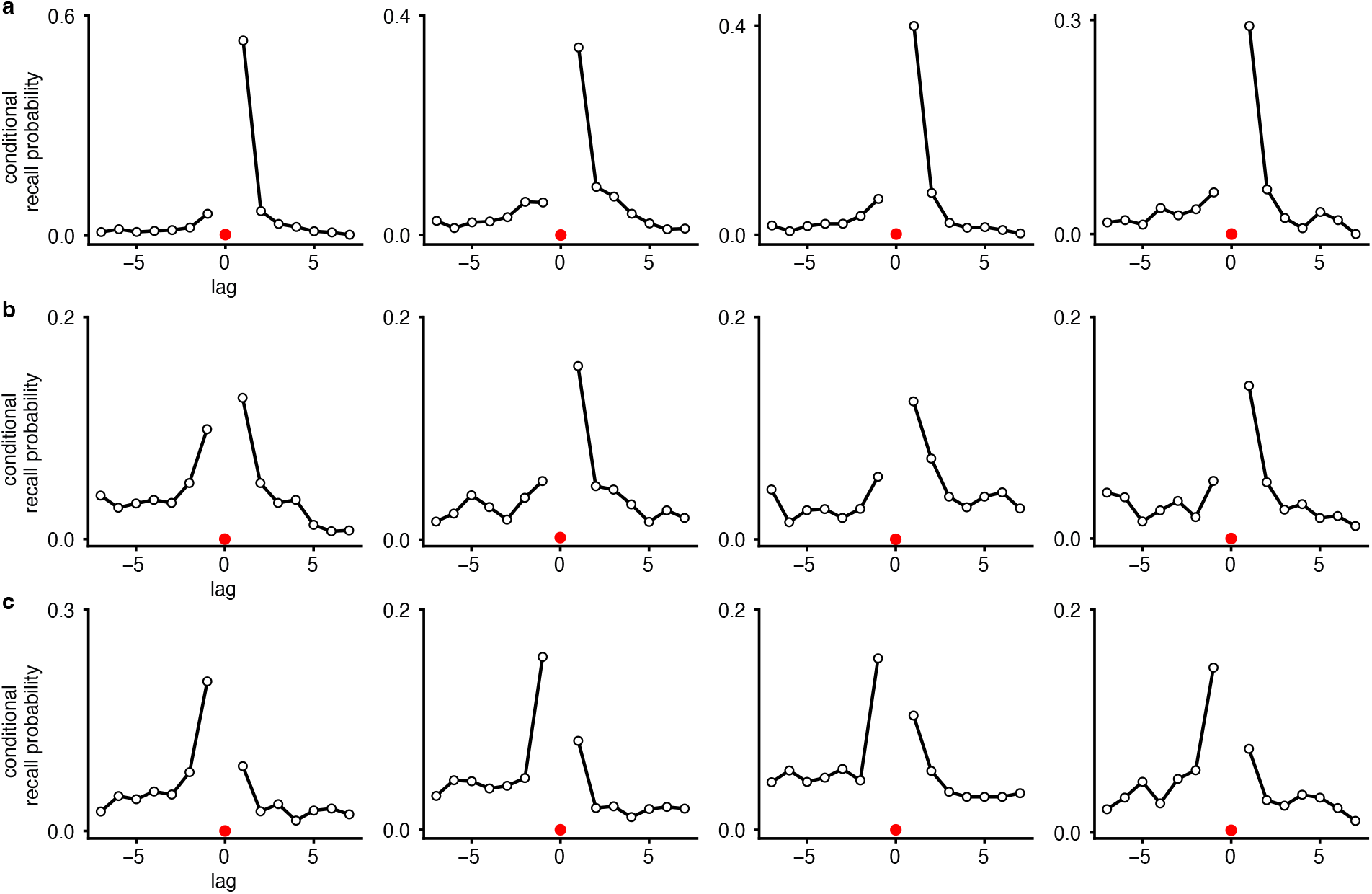
Human behavior recapitulates recall strategies. Four example participants from the PEERS dataset were selected that match the behavior of each strategy. Participants were manually selected through filtering with forward asymmetry and temporal organization score. (a) Conditional recall probability of participants that match the memory palace strategy. (b) Conditional recall probability of participants that match the TCM-like forward strategy. (c) Conditional recall probability of participants that match the TCM-like backward strategy.

**Extended Data Figure 4:**
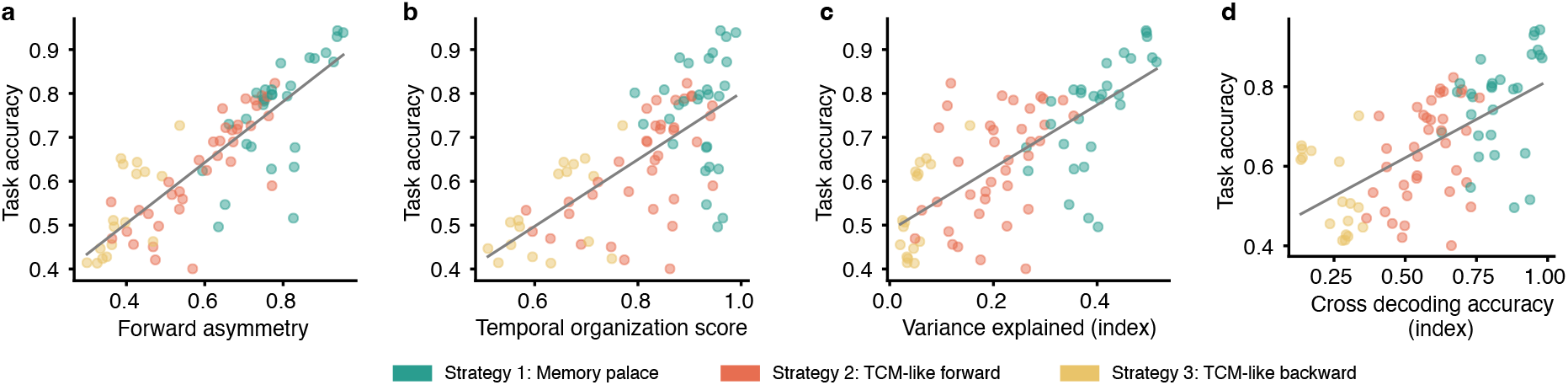
Relationship between strategy metrics and free recall performance. Relationship between (a) Forward asymmetry (Pearson correlation, *r*(81) = 0.71, *p <* 0.0001), (b) temporal organization score (Pearson correlation, *r*(81) = 0.46, *p <* 0.0001), (c) variance explained by index (Pearson correlation, *r*(81) = 0.51, *p <* 0.0001), (d) cross-decoding accuracy of index and the task performance (Pearson correlation, *r*(81) = 0.38, *p <* 0.0001) for 100 models with the same training condition (discount factor = 1.0, % WM flushed = 0.9). Models with free recall performance less than 40% were excluded (n=83). Different colors represent different strategies.

**Extended Data Figure 5:**
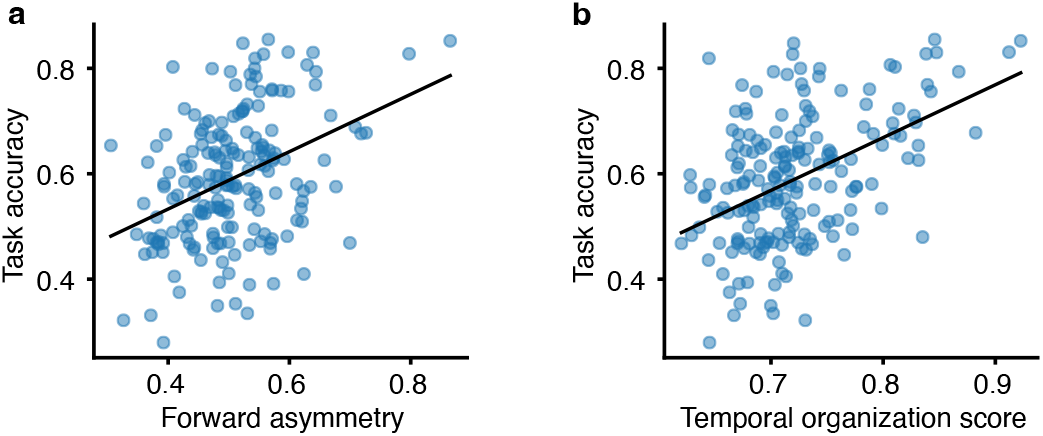
Relationship between behavioral metrics and free recall performance in human data. Relationship between (a) Forward asymmetry (Pearson correlation, *r*(184) = 0.15, *p <* 0.0001) and (b) temporal organization score and the task performance (Pearson correlation, *r*(184) = 0.22, *p <* 0.0001) for 186 participants in the PEERS dataset.

**Extended Data Figure 6:**
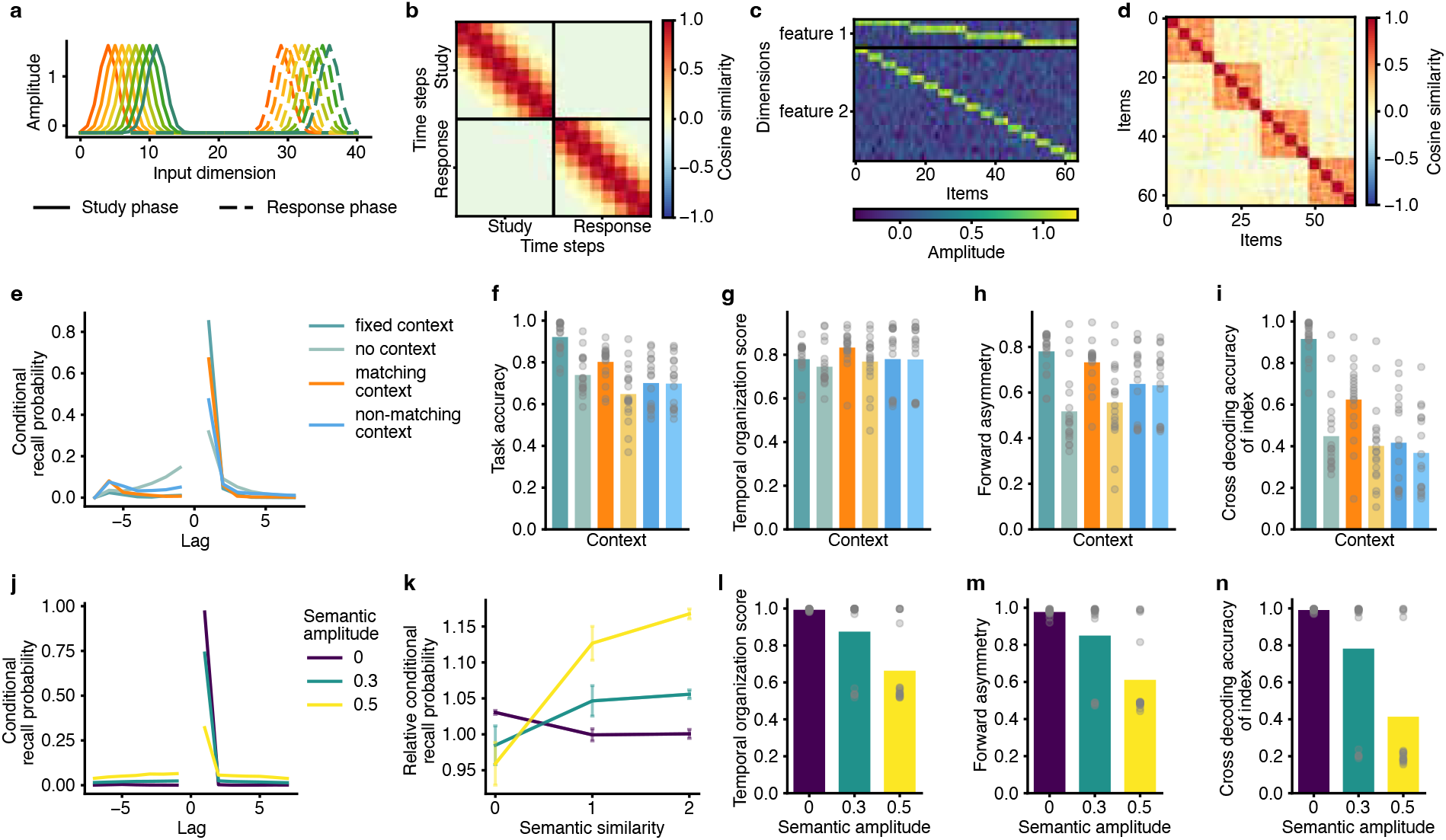
Change of strategy with external and semantic context included as inputs. (a) Example context vectors as inputs when different context is provided in the study phase and in the response phase. Context within a phase drifts slowly, while context across phases are separated. (b) Cosine similarity between contexts provided at different time steps, when context is different in the study phase and in the response phase. Context within a phase has higher similarity when provided closer in time, while context across phases are dissimilar. (c) Semantic vector for each item as inputs. There are 2 features for each item. The first feature is 4 dimensional, and every 16 items share the same feature. The second feature is 16 dimensional, and every 4 items share the same feature. Gaussian noise is added to the vectors. Cosine similarity between the semantic vectors for items. There are 3 levels of semantic similarity. (e) Mean conditional recall probability for models with matching context provided in the study phase and the response phase, and models with unmatched contexts provided across phases. Models with fixed context have the same context vectors across phases and trials, and models with no context have context provided in the study phase but not in the response phase. 20 models with different random seeds were trained for each setting. (f) Mean free recall performance of models trained with the same or different context across phases, and tested with the same or different context, respectively. Dots show individual models for panels (f)-(g) (n=20). (g) Temporal organization score, (h) forward asymmetry, (i) cross-decoding accuracy of index of models trained and tested with the same or different context. (j) Mean conditional recall probability for models trained with different amplitudes of semantic information as inputs. 20 models with different random seeds were trained for each setting. (k) Mean relative conditional recall probability as a function of semantic similarity level for models trained with different amplitudes of semantic information. Error bars denote one standard deviation. (l) Temporal organization score, (m) forward asymmetry, (n) cross-decoding accuracy of index of models trained with different amplitudes of semantic information (n=20).

## References

Anderson, J. R. and Milson, R. (1989). Human memory: An adaptive perspective. Psychological Review, 96(4):703–719.

Basile, A. G. and Toplak, M. E. (2015). Four converging measures of temporal discounting and their relationships with intelligence, executive functions, thinking dispositions, and behavioral outcomes. Frontiers in Psychology, 6:728.

Bennion, K. A., Venturini, M. C., Reyes, H., Cheng, K., Kendra Eng, T., and Antony, J. W. (2025). Classic free recall memory effects using video stimuli. Memory, pages 1–10.

Bernardi, S., Benna, M. K., Rigotti, M., Munuera, J., Fusi, S., and Salzman, C. D. (2020). The geometry of abstraction in the hippocampus and prefrontal cortex. Cell, 183(4):954–967.

Brown, G. D., Preece, T., and Hulme, C. (2000). Oscillator-based memory for serial order. Psychological review, 107(1):127.

Chandra, S., Sharma, S., Chaudhuri, R., and Fiete, I. (2025). Episodic and associative memory from spatial scaffolds in the hippocampus. Nature, 638(8051):739–751.

Cho, K., Van Merriënboer, B., Gulçehre, Ç., Bahdanau, D., Bougares, F., Schwenk, H., and Bengio, Y. (2014). Learning phrase representations using rnn encoder–decoder for statistical machine translation. In Proceedings of the 2014 conference on empirical methods in natural language processing (EMNLP), pages 1724–1734.

Dresler, M., Shirer, W. R., Konrad, B. N., Müller, N. C., Wagner, I. C., Fernández, G., Czisch, M., and Greicius, M. D. (2017). Mnemonic training reshapes brain networks to support superior memory. Neuron, 93(5):1227–1235.

Gershman, S. J., Fiete, I., and Irie, K. (2025). Key-value memory in the brain. Neuron, 113(11):1694–1707.

Glanzer, M. and Cunitz, A. R. (1966). Two storage mechanisms in free recall. Journal of verbal learning and verbal behavior, 5(4):351–360.

Glenberg, A. M., Bradley, M. M., Stevenson, J. A., Kraus, T. A., Tkachuk, M. J., Gretz, A. L., Fish, J. H., and Turpin, B. M. (1980). A two-process account of long-term serial position effects. Journal of Experimental Psychology: Human Learning and Memory, 6(4):355.

Healey, M. K., Crutchley, P., and Kahana, M. J. (2014). Individual differences in memory search and their relation to intelligence. Journal of Experimental Psychology: General, 143(4):1553.

Healey, M. K., Long, N. M., and Kahana, M. J. (2019). Contiguity in episodic memory. Psychonomic bulletin & review, 26(3):699–720.

Hong, M. K., Gunn, J. B., Fazio, L. K., and Polyn, S. M. (2024). The modulation and elimination of temporal organization in free recall. Journal of Experimental Psychology: Learning, Memory, and Cognition, 50(7):1035.

Howard, M. W. and Kahana, M. J. (1999). Contextual variability and serial position effects in free recall. Journal of Experimental Psychology: Learning, Memory, and Cognition, 25(4):923.

Howard, M. W. and Kahana, M. J. (2002). A distributed representation of temporal context. Journal of mathematical psychology, 46(3):269–299.

Jensen, K. T. (2024). An introduction to reinforcement learning for neuroscience. Neurons, Behavior, Data analysis, and Theory.

Johnson, J. D. and Rugg, M. D. (2007). Recollection and the reinstatement of encoding-related cortical activity. Cerebral cortex, 17(11):2507–2515.

Kahana, M. J. (1996). Associative retrieval processes in free recall. Memory & cognition, 24(1):103–109.

Kahana, M. J., Lohnas, L. J., Healey, M. K., Aka, A., Broitman, A. W., Crutchley, P., Crutchley, E., Alm, K. H., Katerman, B. S., Miller, N. E., et al. (2024). The penn electrophysiology of encoding and retrieval study. Journal of Experimental Psychology: Learning, Memory, and Cognition.

Koppenaal, L. and Glanzer, M. (1990). An examination of the continuous distractor task and the “long-term recency effect”. Memory & Cognition, 18(2):183–195.

Li, M. (2026). Code for ‘a neural network model of free recall learns multiple memory strategies’.

Lieder, F. and Griffiths, T. L. (2020). Resource-rational analysis: Understanding human cognition as the optimal use of limited computational resources. Behavioral and brain sciences, 43:e1.

Logan, G. D. and Cox, G. E. (2021). Serial memory: Putting chains and position codes in context. Psychological Review, 128(6):1197.

Lu, Q., Hasson, U., and Norman, K. A. (2022). A neural network model of when to retrieve and encode episodic memories. elife, 11:e74445.

Maguire, E. A., Valentine, E. R., Wilding, J. M., and Kapur, N. (2003). Routes to remembering: the brains behind superior memory. Nature neuroscience, 6(1):90–95.

Mnih, V., Badia, A. P., Mirza, M., Graves, A., Lillicrap, T., Harley, T., Silver, D., and Kavukcuoglu, K. (2016). Asynchronous methods for deep reinforcement learning. In International conference on machine learning, pages 1928–1937. PMLR.

Murdock Jr, B. B. (1962). The serial position effect of free recall. Journal of experimental psychology, 64(5):482.

Neath, I. and Crowder, R. G. (1990). Schedules of presentation and temporal distinctiveness in human memory. Journal of Experimental Psychology: Learning, Memory, and Cognition, 16(2):316.

Polyn, S. M. and Kahana, M. J. (2008). Memory search and the neural representation of context. Trends in cognitive sciences, 12(1):24–30.

Polyn, S. M., Norman, K. A., and Kahana, M. J. (2009a). A context maintenance and retrieval model of organizational processes in free recall. Psychological review, 116(1):129.

Polyn, S. M., Norman, K. A., and Kahana, M. J. (2009b). Task context and organization in free recall. Neuropsychologia, 47(11):2158–2163.

Richards, B. A., Lillicrap, T. P., Beaudoin, P., Bengio, Y., Bogacz, R., Christensen, A., Clopath, C., Costa, R. P., de Berker, A., Ganguli, S., et al. (2019). A deep learning framework for neuroscience. Nature neuroscience, 22(11):1761–1770.

Ritter, S., Wang, J., Kurth-Nelson, Z., Jayakumar, S., Blundell, C., Pascanu, R., and Botvinick, M. (2018). Been there, done that: Meta-learning with episodic recall. In International conference on machine learning, pages 4354–4363. PMLR.

Roediger, H. L. (1980). The effectiveness of four mnemonics in ordering recall. Journal of Experimental Psychology: Human Learning and Memory, 6(5):558.

Romani, S., Katkov, M., and Tsodyks, M. (2016). Practice makes perfect in memory recall. Learning & memory, 23(4):169–173.

Sadeghiyeh, H., Wang, S., Alberhasky, M. R., Kyllo, H. M., Shenhav, A., and Wilson, R. C. (2020). Temporal discounting correlates with directed exploration but not with random exploration. Scientific reports, 10(1):4020.

Sederberg, P. B., Howard, M. W., and Kahana, M. J. (2008). A context-based theory of recency and contiguity in free recall. Psychological review, 115(4):893.

Sederberg, P. B., Miller, J. F., Howard, M. W., and Kahana, M. J. (2010). The temporal contiguity effect predicts episodic memory performance. Memory & cognition, 38:689–699.

Smith, S. M. and Vela, E. (2001). Environmental context-dependent memory: A review and meta-analysis. Psychonomic bulletin & review, 8(2):203–220.

Song, H., Lu, Q., Nguyen, T. T., Chen, J., Leong, Y. C., Rosenberg, M. D., Ching, S., and Zacks, J. M. (2025). A neural network with episodic memory learns causal relationships between narrative events. bioRxiv, pages 2025–09.

Tan, L., Ward, G., Paulauskaite, L., and Markou, M. (2016). Beginning at the beginning: Recall order and the number of words to be recalled. Journal of Experimental Psychology: Learning, Memory, and Cognition, 42(8):1282.

Tulving, E. and Thomson, D. M. (1973). Encoding specificity and retrieval processes in episodic memory. Psychological review, 80(5):352.

Unsworth, N. (2019). Individual differences in long-term memory. Psychological Bulletin, 145(1):79.

Wagner, I. C., Konrad, B. N., Schuster, P., Weisig, S., Repantis, D., Ohla, K., Kühn, S., Fernández, G., Steiger, A., Lamm, C., et al. (2021). Durable memories and efficient neural coding through mnemonic training using the method of loci. Science advances, 7(10):eabc7606.

Wang, J. X., Kurth-Nelson, Z., Kumaran, D., Tirumala, D., Soyer, H., Leibo, J. Z., Hassabis, D., and Botvinick, M. (2018). Prefrontal cortex as a meta-reinforcement learning system. Nature neuroscience, 21(6):860–868.

Whittington, J. C., Muller, T. H., Mark, S., Chen, G., Barry, C., Burgess, N., and Behrens, T. E. (2020). The tolman-eichenbaum machine: unifying space and relational memory through generalization in the hippocampal formation. Cell, 183(5):1249–1263.

Wilding, J. M. and Valentine, E. R. (1997). Superior memory. Psychology Press.

Zaromb, F. M., Howard, M. W., Dolan, E. D., Sirotin, Y. B., Tully, M., Wingfield, A., and Kahana, M. J. (2006). Temporal associations and prior-list intrusions in free recall. Journal of Experimental Psychology: Learning, Memory, and Cognition, 32(4):792.

Zhang, J., Shi, X., King, I., and Yeung, D.-Y. (2017). Dynamic key-value memory networks for knowledge tracing. In Proceedings of the 26th international conference on World Wide Web, pages 765–774.

Zhang, Q., Griffiths, T. L., and Norman, K. A. (2023). Optimal policies for free recall. Psychological Review, 130(4):1104.

