## Supplementary Figures 1-5 for "A neural network model of free recall learns multiple memory strategies"

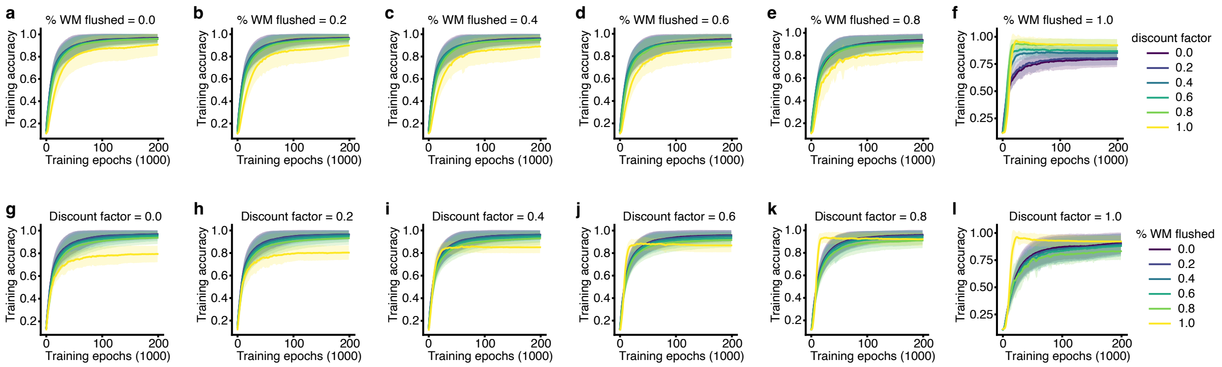


Supplementary Figure 1: **Training curves for all models.** (a-f) Training curves for models with different discount factor under the same percentage of working memory flushed. 20 models with different random seeds were trained for each setting. (g-l) Training curves for models with different percentage of working memory flushed under the same discount factor. 20 models with different random seeds were trained for each setting. Error bands represent the standard deviation across all models with different random seeds.

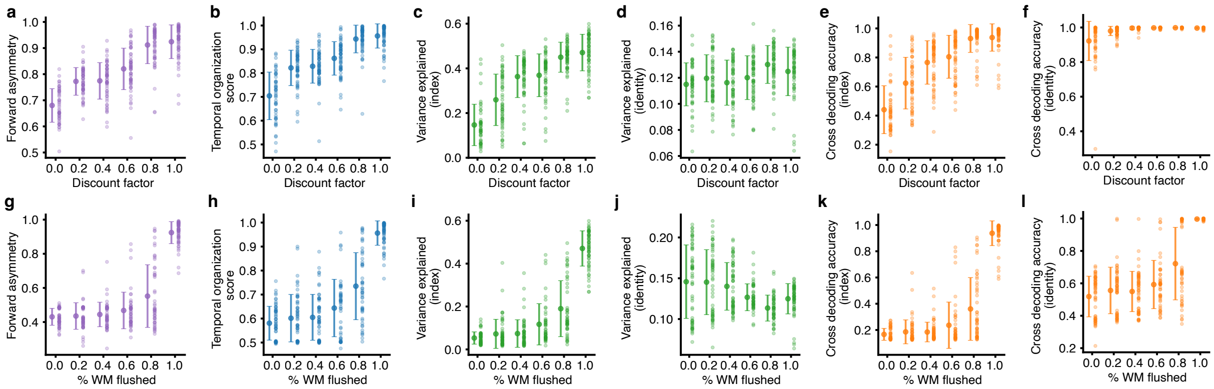


Supplementary Figure 2: **Relationship between strategy metrics with influencing factors of the strategy.** (a) Forward asymmetry, (b) temporal organization score, (c) variance explained by index, (d) variance explained by identity, (e) cross-decoding accuracy of index, (f) cross-decoding accuracy of identity changing with the discount factor. (g-l) The metrics changing with percentage of working memory flushed between the study and response phases. 20 models with different random seeds were trained for each setting. For each hyperparameter setting, the mean and standard deviation were plotted as the error bar, and each model was plotted as a scatter.

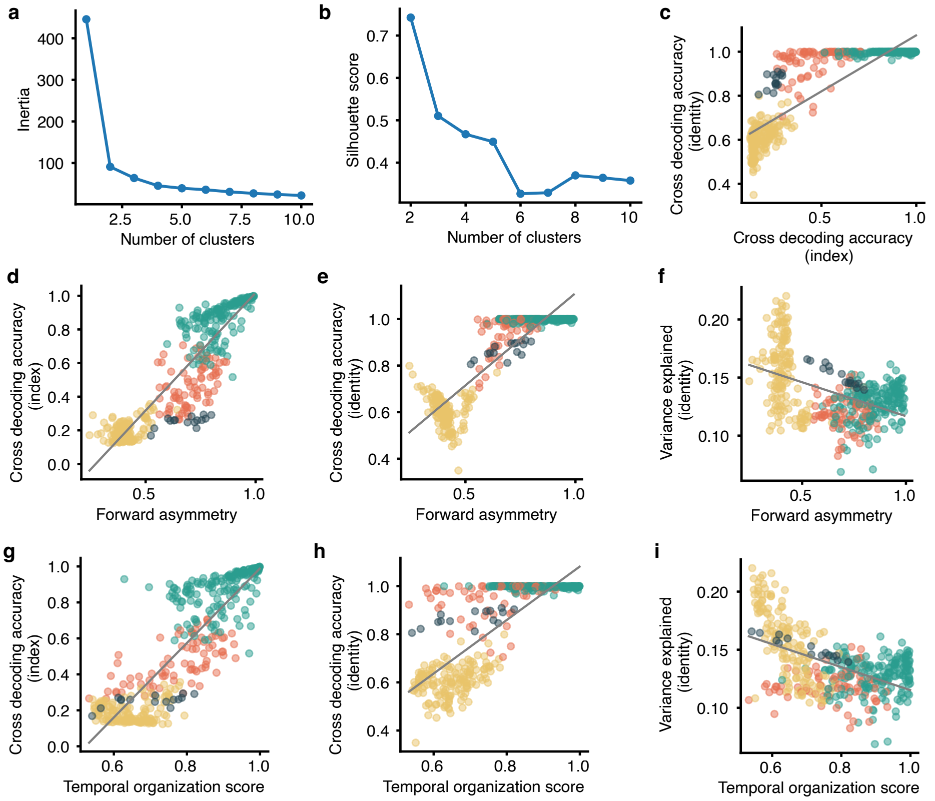


Supplementary Figure 3: **Relationship between metrics used for clustering.** (a) Inertia of clusters in relation with the number of clusters. (b) Sihouette score of different number of clusters. (c) Relation between cross-decoding accuracy of index and identity (Pearson correlation, $r(403) = 0.87, p < 0.0001$). (d) Relation between forward asymmetry and cross-decoding accuracy of index (Pearson correlation, $r(403) = 0.93, p < 0.0001$). (e) Relation between forward asymmetry and cross-decoding accuracy of identity (Pearson correlation, $r(403) = 0.89, p < 0.0001$). (f) Relation between forward asymmetry and explained variance by identity (Pearson correlation, $r(403) = -0.48, p < 0.0001$). (g) Relation between temporal organization score and cross-decoding accuracy of index (Pearson correlation, $r(403) = 0.87, p < 0.0001$). (h) Relation between temporal organization score and cross-decoding accuracy of identity (Pearson correlation, $r(403) = 0.80, p < 0.0001$). (i) Relation between temporal organization score and explained variance by identity (Pearson correlation, $r(403) = -0.51, p < 0.0001$).

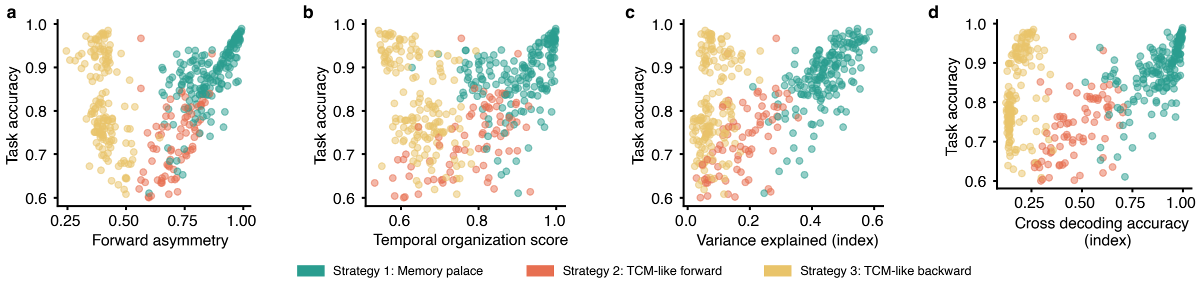


Supplementary Figure 4: **Model performance for all models with different training conditions.** Relationship between (a) Forward asymmetry, (b) temporal organization score, (c) variance explained by index, (d) cross-decoding accuracy of index and the task performance for all models with different training conditions involved in clustering.

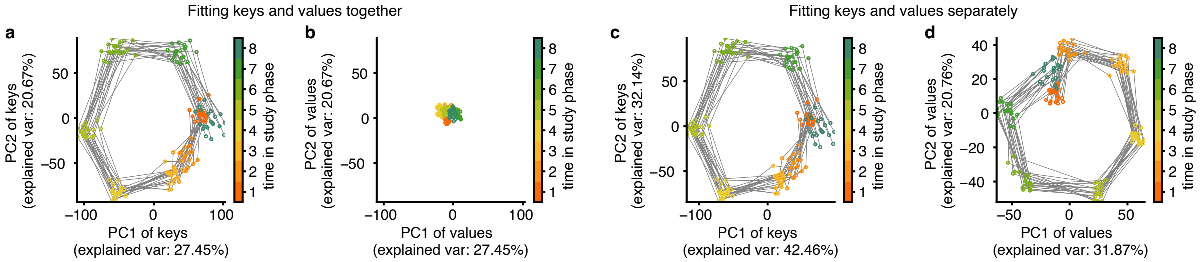


Supplementary Figure 5: **PCA on keys and values for the key-value memory network.** (a, b) PCA trajectories of keys and values for 20 trials when fitting a PCA on concatenated keys and values. (c, d) PCA trajectories of keys and values for 20 trials when fitting them separately. This shows that while both keys and values represented some index information, the representations were not in the same space.
